# Arbitration between insula and temporoparietal junction subserves framing-induced boosts in generosity during social discounting

**DOI:** 10.1101/841338

**Authors:** Manuela Sellitto, Susanne Neufang, Adam Schweda, Bernd Weber, Tobias Kalenscher

**Affiliations:** Comparative Psychology, Institute of Experimental Psychology, Heinrich Heine University Düsseldorf, Universitätsstraße 1, 40225 Düsseldorf, Germany; Department of Psychiatry and Psychotherapy, Medical Faculty, Heinrich Heine University Düsseldorf, Bergische Landstraße 2, 40629 Düsseldorf, Germany; Institute of Experimental Epileptology and Cognition Research, Universitätsklinikum Bonn, Sigmund-Freud Straße 25, 53127 Bonn, Germany

**Keywords:** DCM, fMRI, framing effect, insula, social discounting, TPJ, VMPFC

## Abstract

Generosity toward others declines across the perceived social distance to them. Here, participants chose between selfish and costly generous options in two conditions: in the gain frame, a generous choice yielded a gain to the other; in the loss frame, it entailed preventing the loss of a previous endowment to the other. Social discounting was reduced in the loss compared to the gain frame, implying increased generosity toward strangers. Using neuroimaging tools, we found that while the temporoparietal junction (TPJ) and the ventromedial prefrontal cortex (VMPFC) subserved generosity in the gain frame, the insular cortex was selectively recruited during generous choices in the loss frame. We provide support for a network-model according to which TPJ and insula differentially promote generosity by modulating value signals in the VMPFC in a frame-dependent fashion. These results extend our understanding of the insula role in nudging prosocial behavior in humans.

## Introduction

Most human societies are collaborative. Collaboration offers benefits to their members that they would not be able to achieve individually. However, societies can only function efficiently if their members are willing to contribute to causes whose beneficiaries are abstract and anonymous, such as public goods, and/or to causes whose beneficiaries are socially remote, as it is often the case with wealth redistribution for social welfare, public health insurance, or state pension systems (see also Kalenscher, 2014). Most people are indeed willing to sacrifice own resources for the welfare of others (Nowak, 2006; Rilling & Sanfey, 2011), but their generosity typically declines steeply with social distance between them and the recipients of help, a phenomenon called social discounting (Jones & Rachlin, 2006; Strombach et al., 2015). Hence, while people are ready to provide costly support to friends, relatives, and acquaintances, they are less inclined to help remote strangers.

The social discount function is idiosyncratic (Kalenscher 2017; Vekaria et al. 2017; Archambault, Kalenscher, De Laat 2019; Tiokhin et al. 2019), but it is far from stable within and across individuals. For instance, we and others have shown that participants from individualistic or collectivistic cultures (Strombach et al., 2014) differ in their attitude towards the welfare of socially close peers; that psychosocial stress (Margittai et al., 2015) and neurohormonal stress action (Margittai, Van Wingerden, Schnitzler, Joels, & Kalenscher, 2018) can increase generosity towards socially close friends and acquaintances; and that the level of prosociality towards socially close others depends on gender and cognitive load (Soutschek et al., 2017; Strombach, Margittai, Gorczyca, & Kalenscher, 2016). We further showed that disrupting the temporoparietal junction (TPJ) – a brain region we recently identified as a central hub orchestrating the balance between egocentric and other-regarding preferences in social discounting (Strombach et al., 2015), and which is also associated with perspective taking (Tusche, Bo, Kanske, Trautwein, & Singer, 2016) and theory of mind (Saxe & Kanwisher, 2003) – by means of transcranial magnetic stimulation increases the steepness of social discounting (Soutschek, Ruff, Strombach, Kalenscher, & Tobler, 2016), thus lowering the willingness to support socially remote strangers.

This body of evidence suggests that the degree by which individuals value socially close and distant others’ well-being is highly malleable. However, despite its paramount theoretical and societal significance, means to *increase* the inclination for costly support of socially remote beneficiaries are elusive.

Here, we provide behavioral and neural evidence for a simple manipulation that aims at significantly increasing individuals’ willingness to costly support socially remote others. We make use of the observation that people are more sensitive to others’ losses than gains (Bardsley, 2008; Dreu, 1997; Evans & Beest, 2017; Everett, Faber, & Crockett, 2015; Li et al., 2017; Liu et al., 2020; Schweda, Margittai, & Kalenscher, 2020; Sip, Smith, Porcelli, Kar, & Delgado, 2015; Smith, Sip, & Delgado, 2015; Wang, Rao, & Zheng, 2016; Xiao et al., 2016; Zheng, Wang, & Zhu, 2010), and are consequently strongly reluctant to increase their own payoff at the expense of others’ welfare (Baumeister, Stillwell, & Heatherton, 1994; Chang & Sanfey, 2013; Chang, Smith, Dufwenberg, & Sanfey, 2011; Crockett et al., 2014; List, 2007). We hypothesized that participants would be more altruistic towards others, including socially remote strangers, if a costly generous choice was framed as preventing a monetary loss to others rather than granting them a gain, even if actual economic outcomes were equivalent. In other words, we expected that the way a prosocial decision problem was framed mattered for the shape of the social discount function.

To test this hypothesis, we elicited social preferences in a standard version of the social discounting task (gain frame; Strombach et al., 2015) as well as in a loss frame variant. In each trial, participants decided to share money with other individuals on variable social distance levels. They chose between a selfish option, yielding high own-payoff and zero other-payoff, and a generous option, yielding a lower own-payoff and a non-zero other-payoff. The main difference between conditions was the way the decision problem was described: in the gain frame, a costly generous choice would yield an equivalent gain to the other player, while, in the loss frame, it would imply preventing the loss of a previous endowment to the other player. Importantly, the payoff structure was mathematically identical across frame conditions, i.e., the choice alternatives in the loss frame yielded identical own- and other-payoffs to those in the gain frame. Participants were explicitly instructed that the other persons would only be informed about the final outcome, but not about their endowment, or the loss of it; hence, they knew about the economic equivalence across frames.

We show in a series of independent studies that participants were more reluctant to make a selfish choice if this implied a loss of the endowment to the other, resulting in a substantially flatter social discount function in the loss than the gain frame and, hence, higher generosity towards socially remote others.

To obtain further insights into the psychological and neural mechanisms underlying this framing effect on social discounting, we measured blood oxygen level-dependent (BOLD) responses while participants performed both frame conditions of the social discounting task. We hypothesized that the psychological motives underlying generosity were frame-dependent and dissociable on the neural level. Consistent with our previous work (Strombach et al., 2015), we predicted that generosity in the gain frame was vicariously rewarding and the result of the resolution of the conflict between selfish and altruistic motives. Specifically, generosity in (Strombach et al., 2015) was associated with activity in TPJ, suggesting facilitation in overcoming the egoism bias via the modulation of value signals in the ventromedial prefrontal cortex (VMPFC), a brain structure known to represent own and vicarious reward value (Bartra, Mcguire, & Kable, 2013; Mobbs et al., 2009), through the integration of other-regarding utility. In line with (Soutschek et al., 2016; Strombach et al., 2015), we therefore expected that generous choices in the gain frame would elicit activation of the VMPFC along with TPJ. Conversely, in the loss frame, we expected that the disinclination to maximize own-gain at the expense of other-loss was motivated by the desire to comply to social norms, such as the respect of others’ property rights, or the do-no-harm principle. We therefore hypothesized increased activity in brain regions that are implicated in the social sentiments that motivate individuals to comply to social norms, such as the negative emotions experienced during social norm transgressions, e.g., guilt and shame, as well as the aversive experience of unfairness and inequality (Montague, Lohrenz, & Humphrey, 2007; Xiang, Lohrenz, & Montague, 2013). Such social sentiments have been consistently associated with the insular cortex (Chang & Sanfey, 2013; Chang et al., 2011; Civai, Crescentini, Rustichini, & Rumiati, 2012; Corradi-Dell’Acqua, Civai, Rumiati, & Fink, 2013; Gu et al., 2015; Lallement et al., 2013; Oldham, Murawski, Fornito, Youssef, & Lorenzetti, 2018; Samanez-larkin, Hollon, Carstensen, & Knutson, 2008; Siebenthal et al., 2017; Spitzer, Fischbacher, Herrnberger, Grön, & Fehr, 2007; Tomasino et al., 2013; Wang, Rao, & Zheng, 2017; Yu, Hu, Hu, & Zhou, 2014). Our results support our main hypothesis that frame-dependent choice motives were associated with distinct neural signatures. In the gain frame, we replicate VMPFC and TPJ involvement in generous choice (Hutcherson, Bushong, & Rangel, 2015; Strombach et al., 2015), while in the loss frame, we identify the insular cortex as the core component in processing altruistic motives, prompting generosity by biasing people away from selfish desires if these caused harm to others.

## Results

### Social discounting is flatter in the loss than the gain frame

First, in a set of behavioral experiments, we established that our framing manipulation affected generosity towards socially distant others. In a within-subject design, we elicited social preferences in a standard version of the social discounting task (the gain frame; (Strombach et al., 2015)) as well as in a loss frame variant (see **Fig. 1**), interleaved in a trial-by-trial fashion. In the gain frame, participants played with other persons at variable social distance levels, and made choices between a selfish alternative, yielding higher monetary payoff to the participant and zero payoff to the other, and a generous alternative, always yielding a lower own-payoff of €75 along with a payoff of €75 to the other. In the loss frame, participants were first informed that the other person had received an initial endowment of €75. The selfish alternative yielded a variable, higher own-payoff as well as the loss of the €75 endowment to the other person, hence, resulting in a zero payoff to the other; the generous alternative yielded a fixed €75 payoff to the participant, and no further consequence to the other, thus leaving her with the initial €75 endowment. Crucially, the payoff structure was mathematically equivalent across both frame conditions, i.e., the choice alternatives in the loss frame yielded identical own- and other-payoffs to those in the gain frame. The main difference between conditions was that, in the gain frame, a generous choice would imply a gain of €75 to the other, while in the loss frame, a generous choice would imply preventing the loss of the previous €75 endowment. Importantly, participants were repeatedly instructed that the other person was unaware of her initial endowment, or the loss of it, and that she would only be informed about the final outcome of the payoff after implementing the participant’s choice at the end of the experiment. Task comprehension, in particular regarding participants’ understanding that the other person would only be informed about the final outcome, but not about her endowment, or loss of it, was further stressed during the explanation of the incentivization procedure as well as assessed in post-hoc structured interviews (see Materials and methods). All participants understood the task well.

**Figure 1.**
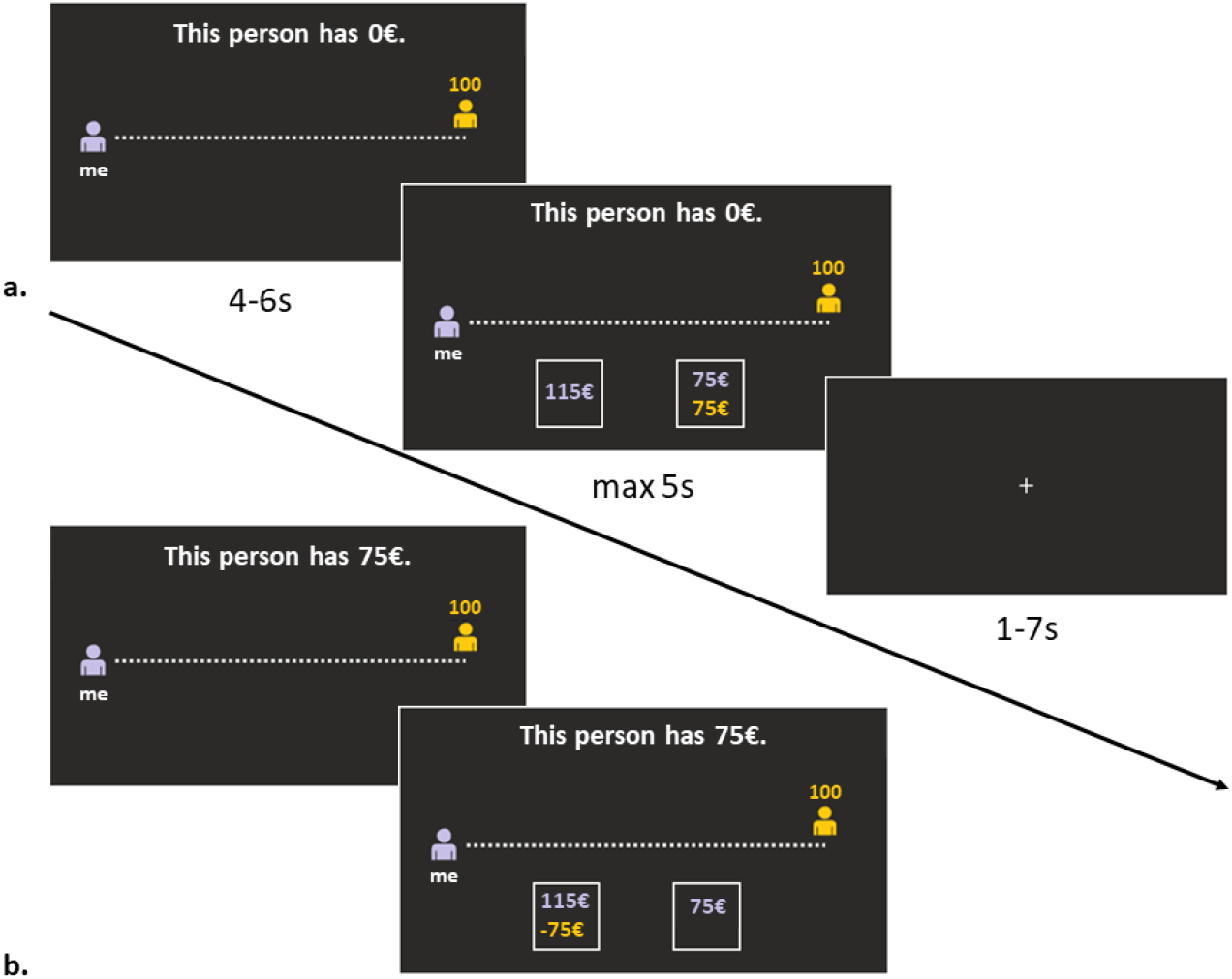
Social discounting task (fMRI study 4). **a.** Trial example of the *gain frame*. **b.** Trial example of the *loss frame*. Each trial started with the presentation of a ruler-based representation of social distance to the other-person, with a left-most purple icon representing the participant and a yellow icon indicating the social distance of the other-person in the current trial (100 in this example). Additionally, participants received information as to the endowment of the other person, i.e., “This person has €0” for the gain frame (**a**), or “This person has €75” for the loss frame (**b**) (4-6s). Afterwards, the two choice options appeared. The selfish alternative was displayed in purple fonts, indicating the own-reward magnitude to the participant (here: €115). Selfish choices implied a null gain for the partner in the gain frame (**a**) or the loss of the initial €75-endowment (in yellow) for the other person in the loss frame (**b**). The generous alternative was displayed in yellow fonts, and always yielded an equal €75 own-reward and €75 other-reward split in the gain frame (**a**), or a €75 own-reward gain and the possibility to keep the €75 other-endowment in the loss frame (**b**). As soon as the two choice options appeared, participants had 5s to choose one of the two alternatives. After a choice was made, or after the 5s were passed, a blank screen with a fixation cross appeared (1-7s), and then a new trial started.

#### Study 1

In a first study, data collection was done online and the task was not incentive-compatible; participants (*n* = 52) were paid a fixed allowance of €8.5. In the social discounting task, participants can either make a selfish choice or a generous choice, in each frame condition. To reconstruct the individual social discount functions, separately for the two frame conditions, we fit a standard hyperbolic model (see Eq. 1 (Jones & Rachlin, 2006; Strombach et al., 2015); Materials and methods) to trial-by-trial binary choices (i.e., either selfish or generous) via a softmax function to estimate the parameter *k*, a measure of the steepness of the social discount function. Additionally, we determined, for each participant and each social distance level, and separately for the two frame conditions, the point at which the participant was indifferent between the selfish and the generous alternative using logistic regression (Strombach et al., 2015). The difference in reward magnitudes for the participant between the two alternatives at the indifference points (see Supplementary results) represented the amount of money a subject was willing to forego (i.e., reward amount foregone) to increase the wealth of another person at a given social distance, and could be construed as a social premium, that is, the price tag participants put on increasing the wealth of the other. We took the estimated parameter V, the intercept at social distance 0, thus the origin of the social discount function (Jones & Rachlin, 2006; Margittai et al., 2015; Margittai et al., 2018; Soutschek et al., 2016; Strombach et al., 2015), as an indicator of generosity towards socially close others (Soutschek et al., 2016; Strombach et al., 2015).

Participants’ generosity dropped much less steeply in the loss compared to the gain frame (median k_gain_ = 0.011 vs. median k_loss_ = 0.005; Wilcoxon matched pairs test: Z = 3.96, p < 0.001; r = 0.56; see supplemental **Fig. S1a**). The difference in social discount functions between frames was most pronounced at high social distance levels, indicating that participants were substantially more generous towards strangers in the loss than the gain frame. We also found a significant difference in V between frame conditions (median V_gain_ = 89 vs. median V_loss_ = 80; Z = 3.32, p < 0.001; r = 0.47) that, however, disappeared when removing all participants with zero discounting from the analysis (see Supplementary results).

These data suggest that participants were strongly more generous towards socially distant others in the loss than the gain frame.

#### Study 2

In a second study, we replicated the results of our first experiment. Data collection was done online and participants (*n* = 30) were reimbursed for their time with a fixed amount of university credits. We again found that participants had flatter social discounting in the loss than the gain frame (median k_gain_ = 0.017 vs. median k_loss_ = 0.008; Z = 2.89, p < 0.004; r = 0.52; see supplemental **Fig. S1b**), and we found no difference in V between frame conditions (median V_gain_ = 80 vs. median V_loss_ = 80; Z = 0.21, p = 0.83). Again, these results held when excluding participants with null discounting.

#### Study 3

Studies 1 and 2 were not incentive-compatible. To determine whether hypothetical versus real payoffs made a difference in the frame effect on social discounting (Vlaev, 2012), we ran a third fully incentive-compatible study in a laboratory setting (*n* = 31). Payoff was contingent on the participants’ choices, and was paid out to self and other, identical to (Strombach et al., 2015) and to the fMRI study 4 (see next paragraph and Materials and methods). Once again, we could replicate the frame effect on *k* (median k_gain_ = 0.021 vs. median k_loss_ = 0.012; Z = 3.16, p < 0.001; r = 0.57; see supplemental **Fig. S1c**), and there was no difference in the V parameter between frame conditions (median V_gain_ = 81 vs. median V_loss_ = 80; Z = 1.43, p = 0.15). These results held when excluding participants with null discounting. Additionally, social desirability, as measured via the Social Desirability Scale (SDS-17; (Stöber, 2001)), did neither explain the V nor the *k* parameters across frame conditions in any of the three studies (see Supplemental materials).

Overall these results suggest that, while generosity to socially close others was comparable between frame conditions, it decayed significantly less steeply across social distance in the loss than in the gain frame, indicating that participants were considerably more generous towards socially distant others in the loss frame.

### Neural mechanisms underlying the frame effect on social discounting

To obtain more substantial insights into the psychological and neural mechanisms underlying this framing effect on social discounting, in study 4, we measured BOLD responses while participants performed both frame variants of the social discounting task. The fundamental premise of our study is that the decision motives and their neural correlates differ between gain and loss frames. Specifically, we reasoned that generosity in the gain frame was mainly stimulated by other-regarding considerations. Conversely, we predicted that generous decisions in the loss frame were motivated by the desire to comply to social norms, such as the *do-no-harm* principle, or the respect of others’ property rights (Somanathan, 2016), infringements of which are associated with negative social sentiments of guilt and shame. To test this idea, we focused on two main hypotheses. We, first, expected that generosity in the gain frame recruited a network of structures, including VMPFC and TPJ (Hutcherson et al., 2015; Strombach et al., 2015), known to represent vicarious reward value and overcoming egoism bias. Second, we hypothesized that brain areas implicated in negative social sentiments of social norm transgressions, such as the insular cortex (e.g. Paulus et al. 2003; Chang et al. 2011; Chang and Sanfey 2013; Lallement et al. 2013; Gu et al. 2015; Seara-cardoso et al. 2016; Sethi and Somanathan 2016; Siebenthal et al. 2017; Wang et al. 2017; Huggins et al. 2018), would be selectively recruited during generous choices in the loss, but not the gain frame.

We first replicated, once more, the behavioral framing effect on social discounting (*n* = 30; two participants were excluded from the behavioral analyses due to bad curve fitting; see Materials and methods). As before, the drop in generosity across social distance was pronouncedly flatter in the loss than the gain frame (median k_gain_ = 0.019 vs. median k_loss_ = 0.009; Wilcoxon matched pairs test: Z = 3.46, p < 0.001; r = 0.65; **Fig. 2**), but, again, there was no difference in the V parameter between conditions (median V_gain_ = 80 vs. median V_loss_ = 80; Z = 1.00, p > 0.31; the results remained identical when excluding participants with null discounting). Additionally, neither social desirability (SDS-17; Stöber 2001), nor perspective taking, empathic concern, personal distress, or fantasy (as measured via the Interpersonal Reactivity Index; IRI; (Davis, 1983; Paulus, 2009) explained the V or the *k* parameters across frame conditions (see Supplemental materials).

**Figure 2.**
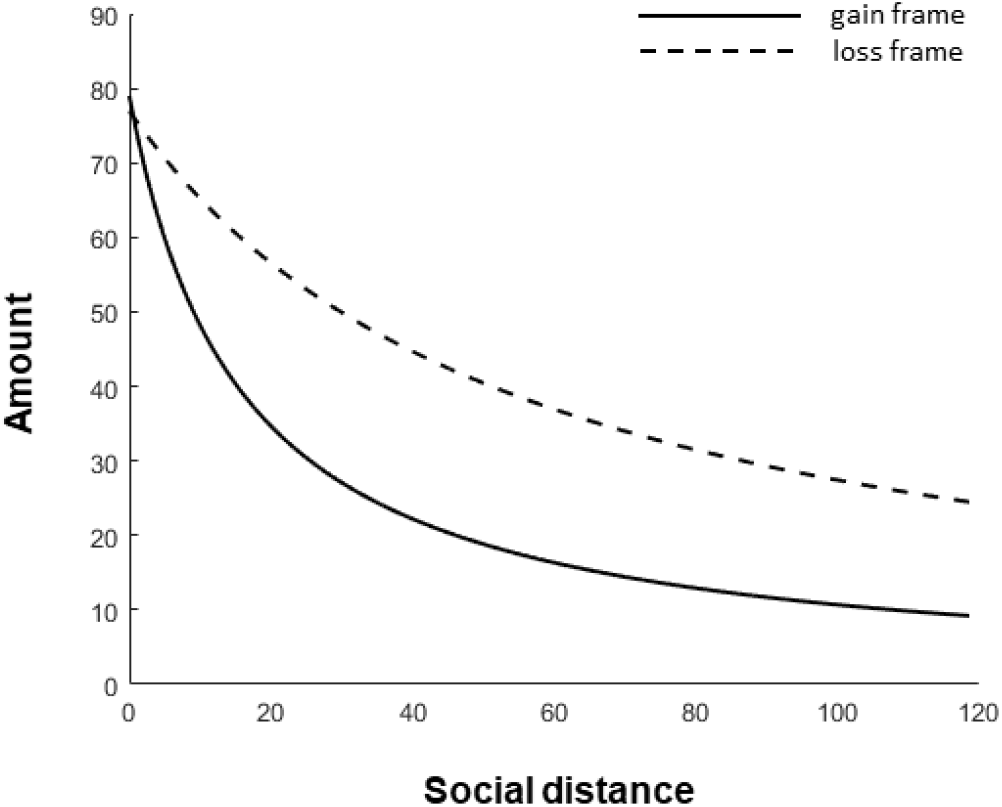
Hyperbolic discounting function (fMRI study 4). The change in generosity across social distances, measured as the social-distance-dependent reward amount that participants were willing to pay to increase the wealth of another person, was captured by a hyperbolic discount model (see main text for details). The figure shows the mean best-fitting hyperbolic functions, averaged across participants, computed separately for the gain frame and the loss frame. The social discounting curve for the loss frame (**dashed line**) was significantly flatter than the social discounting curve for the gain frame (**solid line**).

To test our first hypothesis, we aimed to replicate our previous finding of brain structures known to represent vicarious reward value and overcoming egoism bias in the gain frame (A. Soutschek et al., 2016; Strombach et al., 2015). Our results (**GLM1**; see Materials and methods) indeed revealed clusters located in VMPFC (0, 54, 14, whole-brain p_FWE-corr_ < 0.001) as well as right TPJ (rTPJ; 50, −66, 36, whole-brain p_FWE-corr_ < 0.035) to be selectively activated, in addition to other prefrontal regions, when participants made generous choices in the gain frame relative to generous choices in the loss frame. ROI analyses confirmed significant clusters of activation in both VMPFC (p_FWE-corr_ < 0.000) and rTPJ (p_FWE-corr_ = 0.01). Thus, consistent with (Hutcherson et al., 2015; Strombach et al., 2015), a network comprising VMPFC and rTPJ seems to underlie the motivation for costly generosity in the gain frame (**Fig. 3**; see supplemental **Table S1**). Additionally, selfish amount magnitude, included as trial-by-trial regressor, did not parametrically modulate activity in VMPFC and rTPJ (GLM1; see Materials and methods).

**Figure 3.**
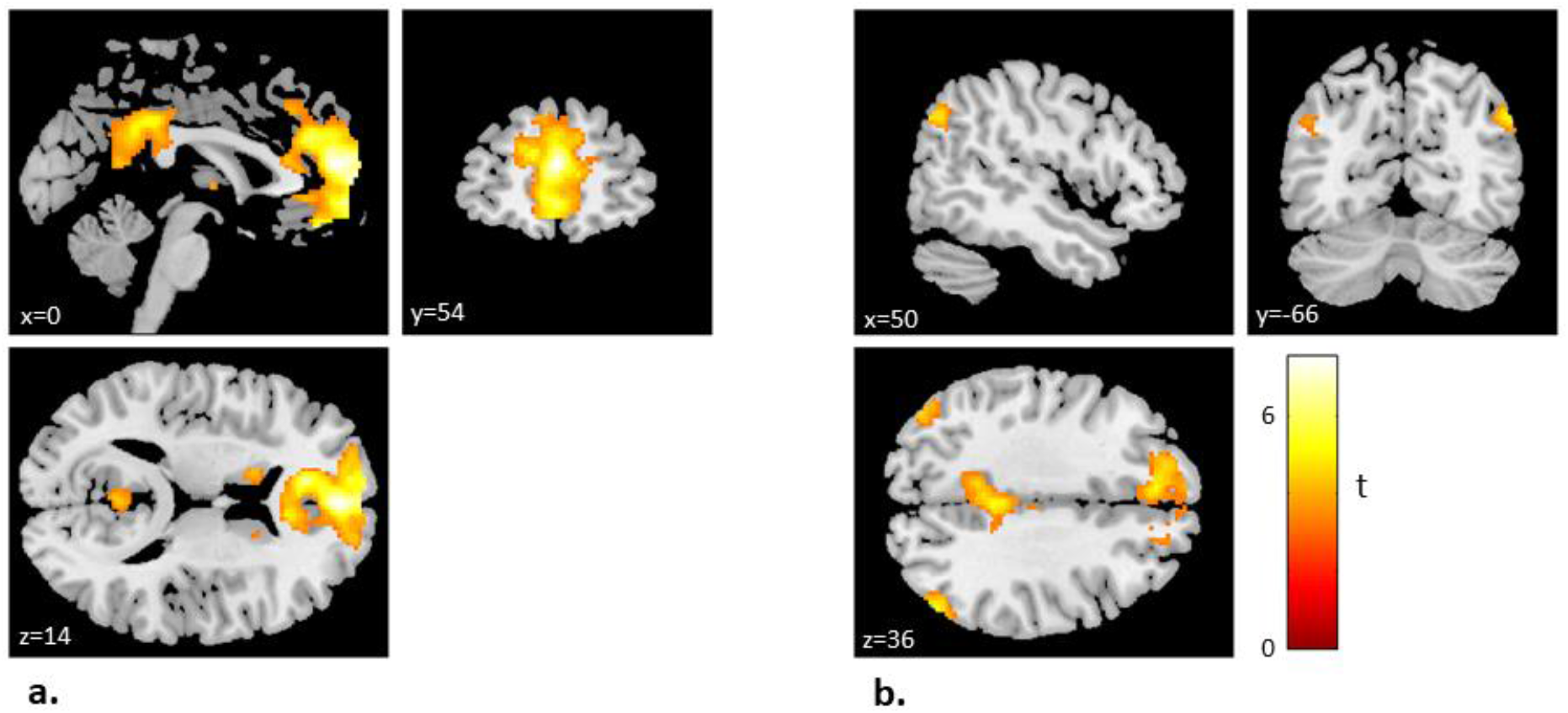
Generous choices in the gain frame correlate with VMPFC and rTPJ activity. VMPFC (MNI peak [0, 54, 14]) (**a**) as well as right TPJ [50, −66, 36] (**b**) were selectively activated during [generous choice in gain frame > generous choice in loss frame] (GLM1; p < 0.05 FWE whole-brain corrected at the cluster level; for illustration purposes, activations are displayed at p < 0.001, uncorrected, minimum cluster size ≥ 5). Color bar indicates T-value.

Our second hypothesis predicted that generosity in the loss frame was motivated by social norm compliance rather than other-regarding considerations; generosity should, consequently, go along with a different neural activation pattern in the loss than the gain frame. In a first step, we attempted to isolate frame-dependent neural correlates, independent of participants’ choices. To this end, we searched for differential neural activity at trial onset, i.e., when participants learned about the social distance level of the other person and which frame was relevant in the current trial (see **Fig. 1**), by contrasting neural activity between the two frames (**GLM2**; see Materials and methods). We found significant activation in the right posterior insula (34, −16, 8, whole-brain p_FWE-corr_ = 0.007) in the loss vs. gain frame contrast, which was accompanied by significant activations in frontal regions, including VMPFC (2, 50, −8, whole-brain p_FWE-corr_ = 0.001), as well as temporal regions (**Fig. 4**; see supplemental **Table S2** for a complete list of activations). ROI analyses confirmed significant clusters of activation in the right insula (p_FWE-corr_ = 0.03) as well as in VMPFC (p_FWE-corr_ = 0.02). The opposite contrast, gain frame vs. loss frame, did not reveal any significant activation. Social distance information, included as trial-by-trial regressor, did not parametrically modulate neural activity in any of these contrasts (GLM2; see Materials and methods), suggesting that the activations in insula and VMPFC reflected frame- but not social distance information.

**Figure 4.**
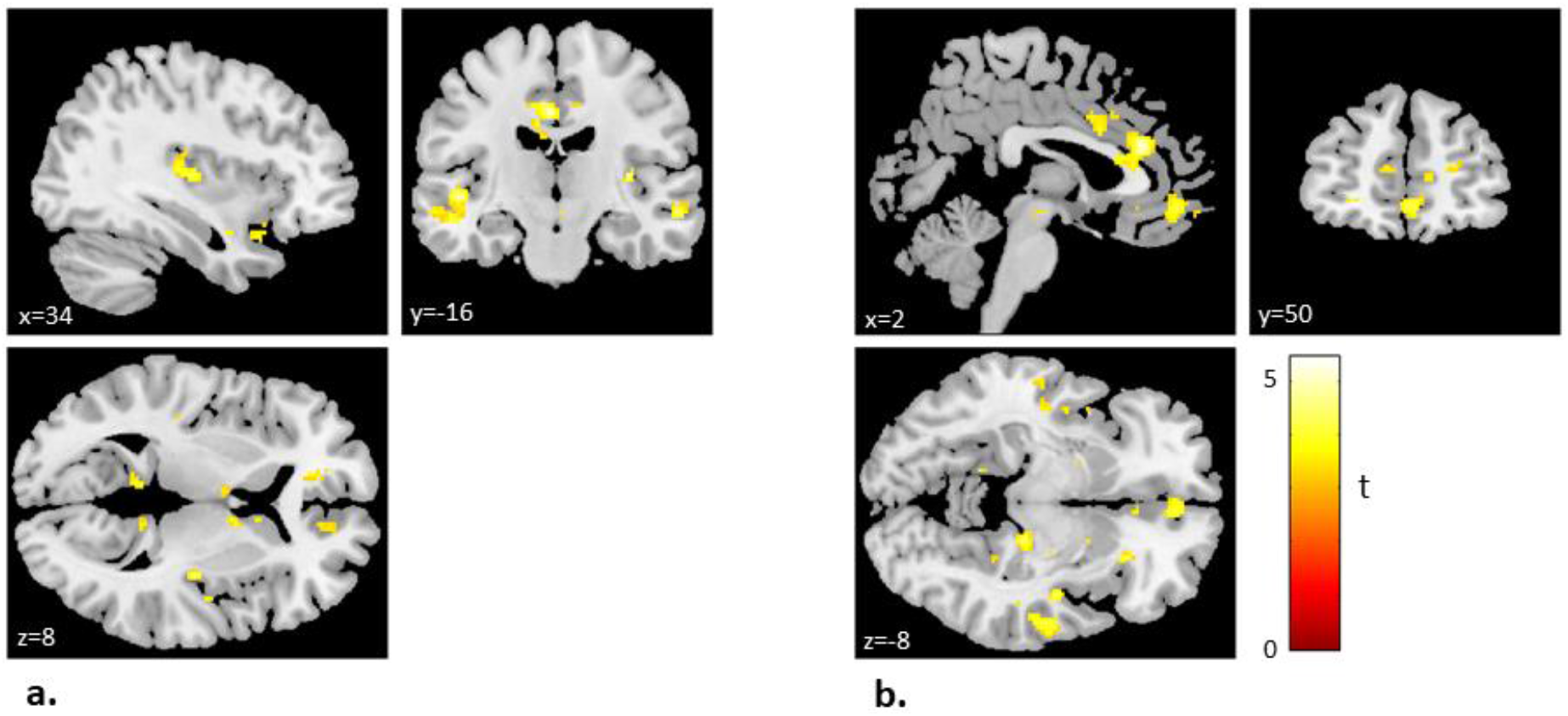
The loss frame information recruits the insula and VMPFC. Insula [34, −16, 8] (**a**) as well as VMPFC [2, 50, −8] (**b**) were selectively activated during [loss frame > gain frame] onset. (GLM2; p < 0.05 ROI FWE-corrected at the cluster level; for illustration purposes, activations are displayed at p < 0.001, uncorrected, minimum cluster size ≥ 5). Color bar indicates T-value.

In support of this conclusion, we found that right anterior insula (42, 4, −4, ROI analysis, p_FWE-corr_ < 0.02; **GLM1**, see Materials and methods), was selectively activated during generous choices in the loss frame relative to generous choices in the gain frame (**Fig. 5**; see supplemental **Table S1**). The location within the insula mask was slightly anterior to the peak activation we found at trial onset.

**Figure 5.**
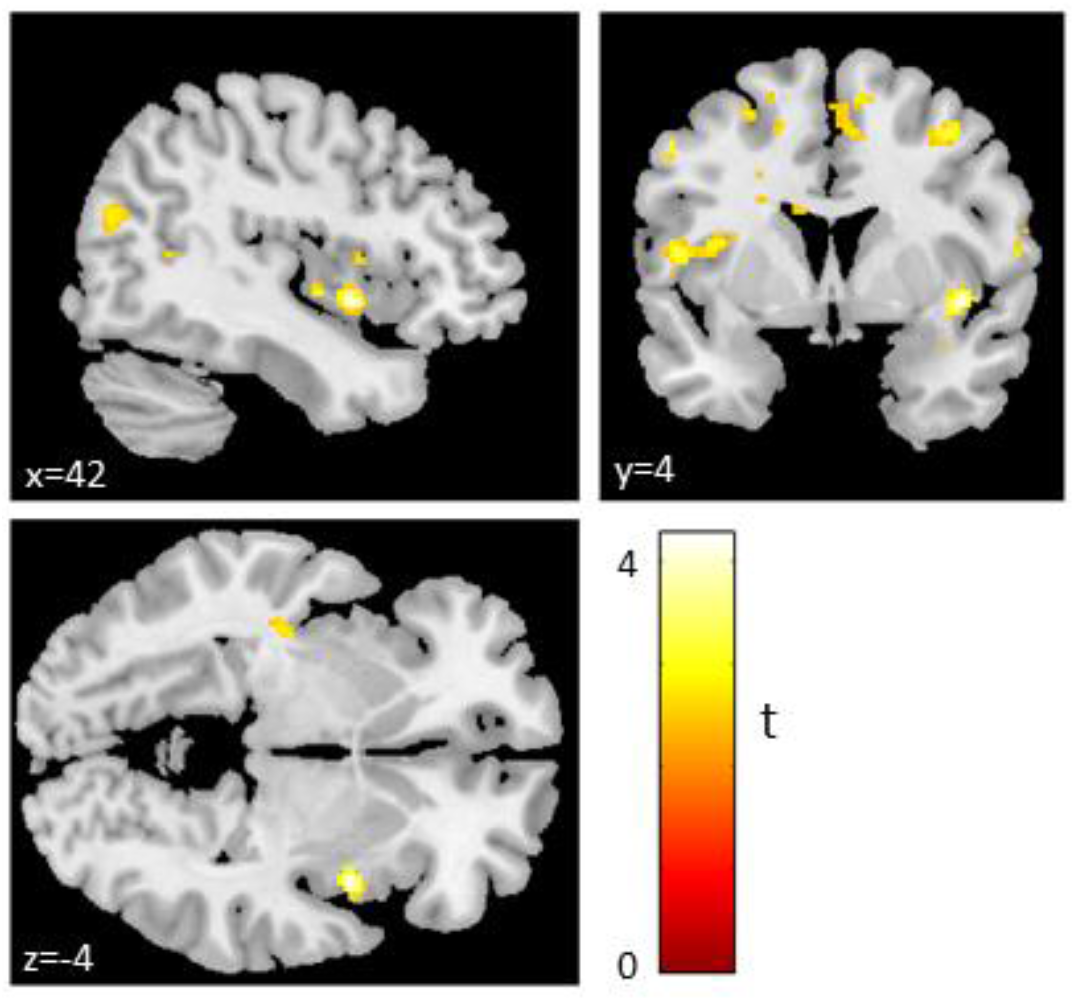
Insula activation underlies generous choice in the loss frame. Insula [42, 4, −4] was selectively activated during [generous choice in loss frame > generous choice in gain frame] (GLM1; p < 0.05 ROI FWE-corrected at the cluster level; for illustration purposes, activations are displayed at p < 0.01, uncorrected, minimum cluster size ≥ 5). Color bar indicates T-value.

To isolate the specific functional contributions of these anterior and posterior clusters within insula, we quantified the extent at which both insula spots correlated with the individual propensity to make generous choices in the gain and the loss frame. To this end, we extracted the parameter estimates at the individual level from an ROI with a seed in both insula clusters (coordinates for the right posterior insula obtained from the results of GLM2, see above [34, −16, 8], and for the anterior insula obtained from the results of GLM1, see above [42, 4, −4]). Parameter estimates were extracted separately for the gain and the loss frame, after pooling all choices (i.e., generous and selfish) within each frame (**GLM3**; see Materials and methods). We then correlated, for each frame separately, the extracted beta estimates with the percentage of generous choices in the respective frame. Both insula activations positively and significantly covaried with generous choices in the loss frame (posterior insula: r = 0.37, p = 0.02; anterior insula: r = 0.33, p = 0.03, one-tailed, medium effect size; **Fig. 6a**), but not in the gain frame (posterior insula: r = 0.07, p = 0.36; anterior insula: r = 0.09, p = 0.31, one-tailed; **Fig. 6b**), thus corroborating the idea that the insula played a different role in promoting generosity in the loss than in the gain frame.

**Figure 6.**
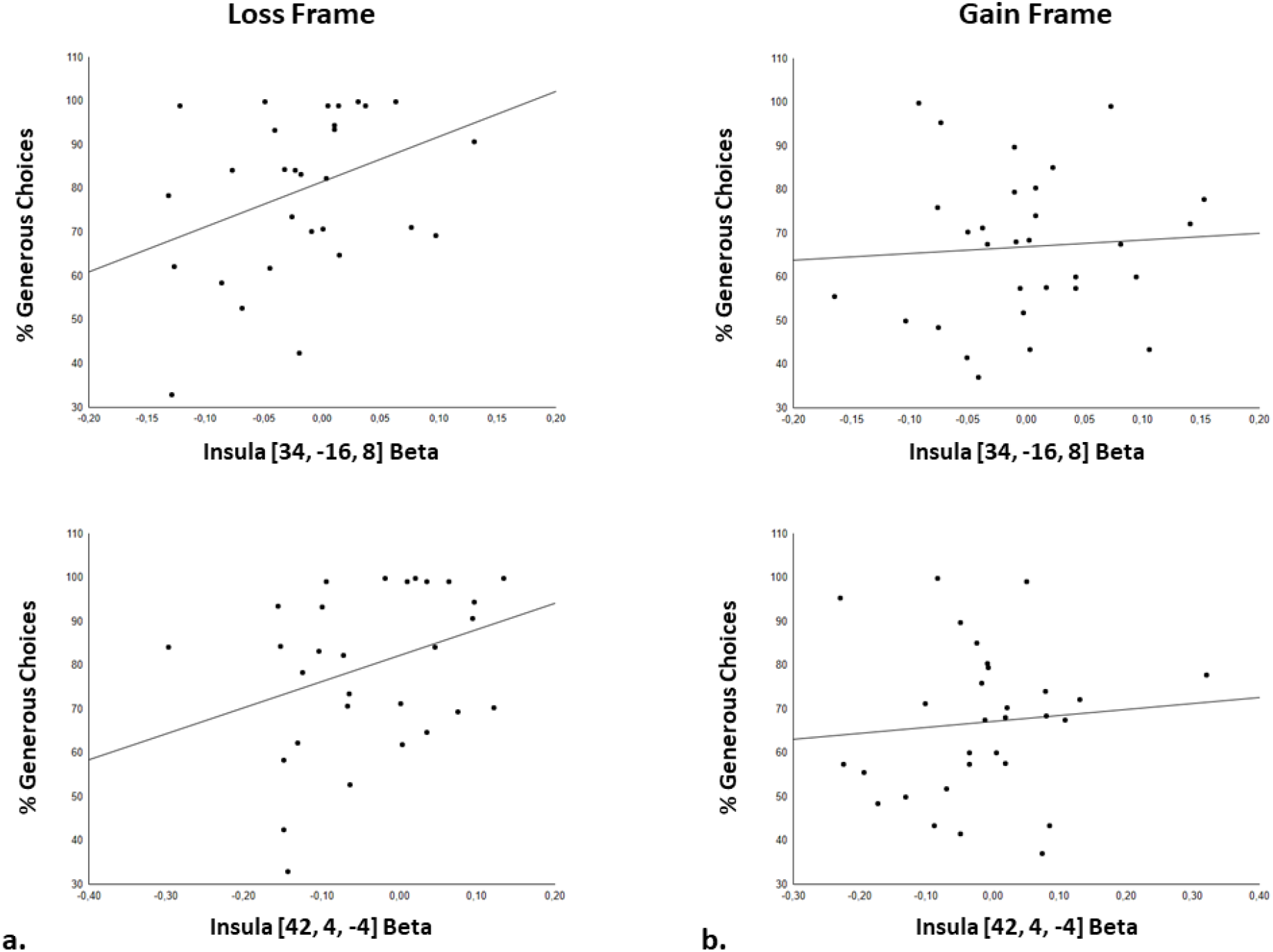
Insula activation correlates with generous choice in the loss frame only. Parameter estimates of insula activation, [34, −16, 8] and [42, 4, −4], were extracted separately for the gain and the loss frame, after pooling all choices (i.e., both generous and selfish) within each frame (GLM3). Insula activation correlated significantly with the percentage of generous choices made in the loss frame (**a**) but not in the gain frame (**b**).

Our analysis so far suggests that insula activation reflects the psychological motives underlying generous choice in the loss frame. However, other explanations of our insula finding are conceivable, too. For instance, participants made more generous choices overall in the loss than the gain frame; i.e., they forewent more own-payoff in the loss than the gain frame, and insula activation might reflect the higher level of reward foregone in the loss frame. Yet, the trial-by-trial regressor of reward amount foregone (GLM1; see Materials and methods) revealed no parametric modulation of insula activity, nor of activity in any other brain region, during generous choices in either frame condition. Additionally, insula activity is unlikely to reflect the own-reward component of the generous alternative because it was fixed (always €75) and, thus, invariant across trials in both frames.

### Frame-dependent modulation of VMPFC activation by rTPJ and insula

We previously provided empirical support for a network model according to which, in a task similar to our gain frame condition, TPJ would facilitate generous decision-making by modulating basic reward signals in the VMPFC, incorporating other-regarding preferences into an otherwise exclusive own-reward value representation, thus computing the vicarious value of a reward to others (Strombach et al., 2015). Here, we expand on this idea and propose that, in addition to the TPJ-VMPFC connectivity in the gain frame, frame-related information in the loss frame would activate insula, which in turn would down-regulate own-value representations in VMPFC, thus promoting generous choices by decreasing the attractiveness of own-rewards. Hence, in brief, we predicted a complex, frame-dependent pattern of connectivity between insula, TPJ, and VMPFC that reflects the different motives underlying generosity in the gain and the loss frame.

To identify the relations between those regions, we estimated their effective connectivity via dynamic causal modeling (DCM) (Friston, Harrison, & Penny, 2003). More specifically, we tested the idea that the frame information at the beginning of each trial would drive increased insula activation selectively in the loss frame, and increased TPJ activation selectively in the gain frame. Additionally, we expected increased endogenous connectivity as well as condition-specific modulation between each respective region with VMPFC. Note that we focused our DCM analysis on the posterior insula cluster only, as we were interested in a baseline frame activation; including the anterior insula cluster, specific for generous choice within the loss frame (see above), might have biased the results in favor of our hypotheses.

In total we defined 15 models (see **Fig. 7**), grouped into two model families: A, which assumed both condition-specific driving inputs and condition-specific modulatory inputs; B, which assumed only condition-specific driving inputs.

**Figure 7.**
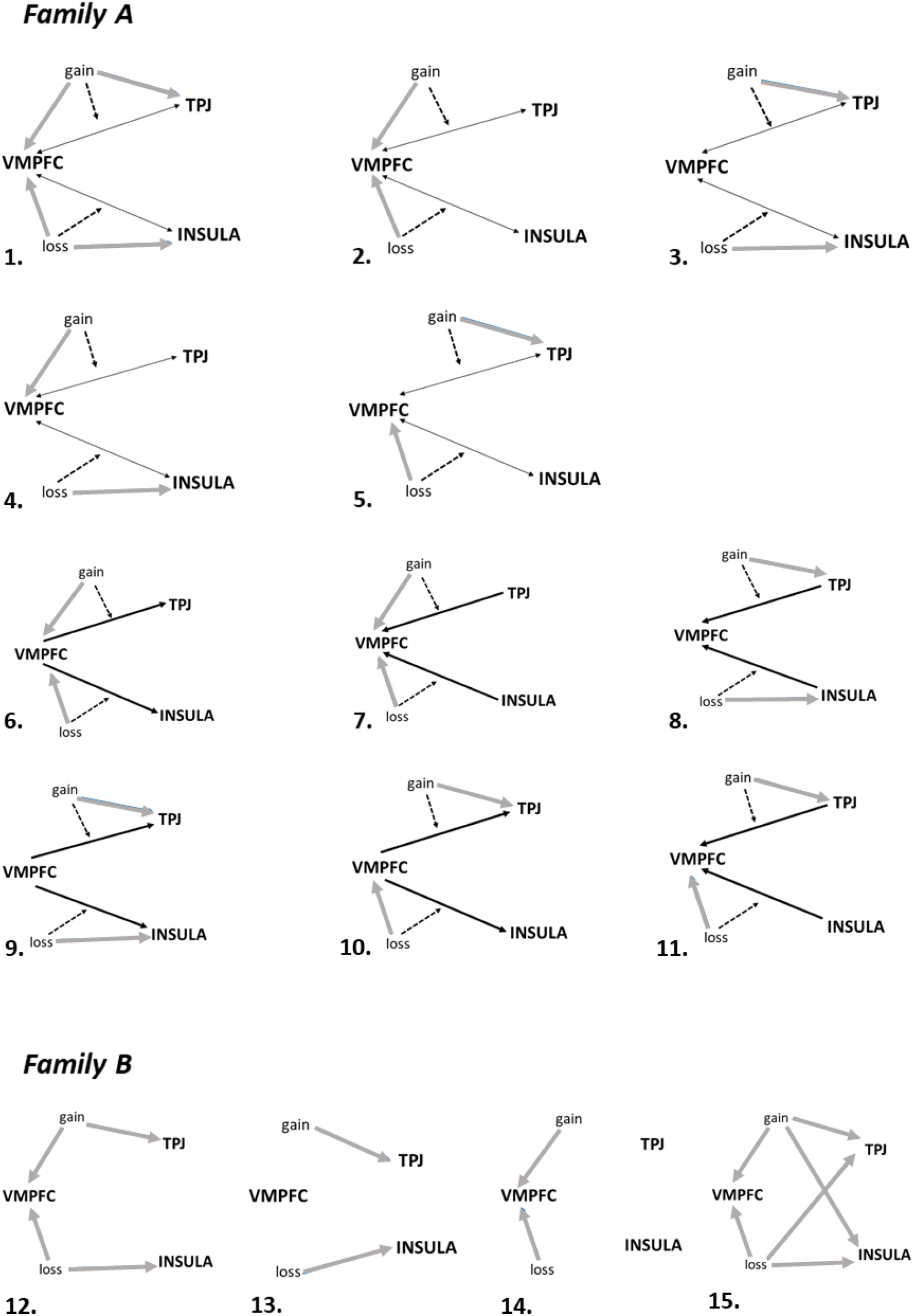
DCM models. Fifteen models were hypothesized to describe the data. Grey lines represent driving inputs. Dashed black lines represent modulatory inputs. Thin black lines represent bidirectional connectivity. Thick black lines represent unidirectional connectivity. Since endogenous connectivity is always assumed between all three regions in all models, it is not represented here. Family A, which assumed both condition-specific driving inputs and condition-specific modulatory inputs, includes models 1 to 11. Family B, which assumed only condition-specific driving inputs, includes models 12 to 15.

Among the two model families tested, model comparison favored family A, i.e., the family of models that assumed condition-specific effects at the level of both driving input and modulatory input (see supplemental **Fig. S2a**). The winning model was model number *5* (sum of the log-evidence SF = −4.0786e+05, exceedance probability xp = 0.6508) (see supplemental **Fig. S2b**), which assumed that the gain frame had an effect on the TPJ and its connectivity with the VMPFC, while the loss frame had an effect on the VMPFC and its connectivity with the insula (i.e., connectivity between regions is assumed to be bidirectional). Concerning the driving inputs, we compared the average activity in TPJ in the gain frame against 0, and the average activity in VMPFC in the loss frame against 0 (we checked, beforehand, that no effect of repetition across runs was present; all ps > 0.18), but none of the driving inputs was significantly different from 0 (all ps > 0.26; **Table 1**).

**Table 1.** DCM estimated parameters of the winning model and statistics. Values are expressed as mean ± standard error (s.e.). Statistics refer to paired t-tests between the modulatory activity and the respective endogenous connectivity, and to one-sample t-tests against 0 for driving inputs. t = t-value; p = p-value; subscript numbers are degrees of freedom; * = p < 0.05. Endo = endogenous connectivity; mod = modulatory connectivity; drivInp = driving input; Gain = gain frame; Loss = loss frame; TPJ = temporoparietal junction; INS = insula; VMPFC = ventromedial prefrontal cortex. Arrows indicate connectivity direction.

| DCM estimated parameters | mean ± s.e. | Statistics |
| --- | --- | --- |
| *endo: TPJ → VMPFC* | 0.05 ± 0.03 | - |
| *endo: VMPFC → TPJ* | 0.02 ± 0.03 | - |
| *endo: INS → VPMFC* | 0.05 ± 0.02 | - |
| *endo: VMPFC → INS* | 0.01 ± 0.02 | - |
| *mod_Gain: TPJ → VMPFC* | 0.03 ± 0.09 | $t_{29} = -0.13$, $p = 0.90$ |
| *mod_Gain: VMPFC → TPJ* | -0.03 ± 0.08 | $t_{29} = -0.53$, $p = 0.60$ |
| *mod_Loss: INS → VMPFC* | -0.22 ± 0.09 | $t_{29} = -2.56$, $p = 0.02^*$ |
| *mod_Loss: VMPFC → INS* | -0.02 ± 0.07 | $t_{29} = -0.30$, $p = 0.80$ |
| *drivInp_Gain: TPJ* | 0.04 ± 0.03 | $t_{29} = 1.31$, $p = 0.20$ |
| *drivInp_Loss: VMPFC* | 0.00 ± 0.03 | $t_{29} = 0.10$, $p = 0.92$ |

Next, when addressing the modulatory inputs, the only significant difference was found in the loss frame for modulatory activity from the insula to VMPFC against the endogenous connectivity from the insula to VMPFC (M_modulatory_ = −0.2158 vs. M_endogenous_ = 0.04672, p = 0.016, Bonferroni corrected), reflecting a significant modulation of endogenous connectivity by the loss frame information (all other ps > 0.60; **Table 1**). In addition, the modulatory input was negative, hinting towards an inhibitory influence of insula on VMPFC in the loss frame (as before, there was no effect of repetition across runs in neither modulatory activity nor endogenous connectivity; all ps > 0.13).

### The mediating role of the insula in the frame effect on social discounting

To provide further support for our idea that the frame effect on social discounting was brought about by a condition-specific neural activity pattern in the insula and VMPFC, and TPJ and VMPFC, respectively, we ran mediation analyses on the relation between frame information, social discounting behaviour, and neural activation in these regions. More specifically, frame was entered as independent variable X (gain and loss), the hyperbolic *k* parameter (gain frame and loss frame) was entered as dependent variable Y, and the neural activations were entered as mediators. We first focused on a model (*model 1*) where neural activations across both frames were treated as parallel mediators. Neural activations included the posterior insula and the anterior insula, TPJ, and VMPFC (both clusters in GLM1 and GLM2) (see Materials and methods for details). Frame condition significantly correlated with all neural activations (all ps < 0.05) with the exception of VMPFC (GLM2) (p = 0.08). Additionally, while the direct effect of the frame condition on *k* was not significant (p = 0.75), its total indirect effect was significant and negative, indicating that all mediators significantly mediated the relation between frame condition and the *k* parameter, and that the effect was stronger on the loss than the gain frame (partially standardized B = −0.41, SE = 0.21, 95% biased-corrected confidence interval (CI) −0.88 to −0.06) (**Fig. 8a**).

**Figure 8.**
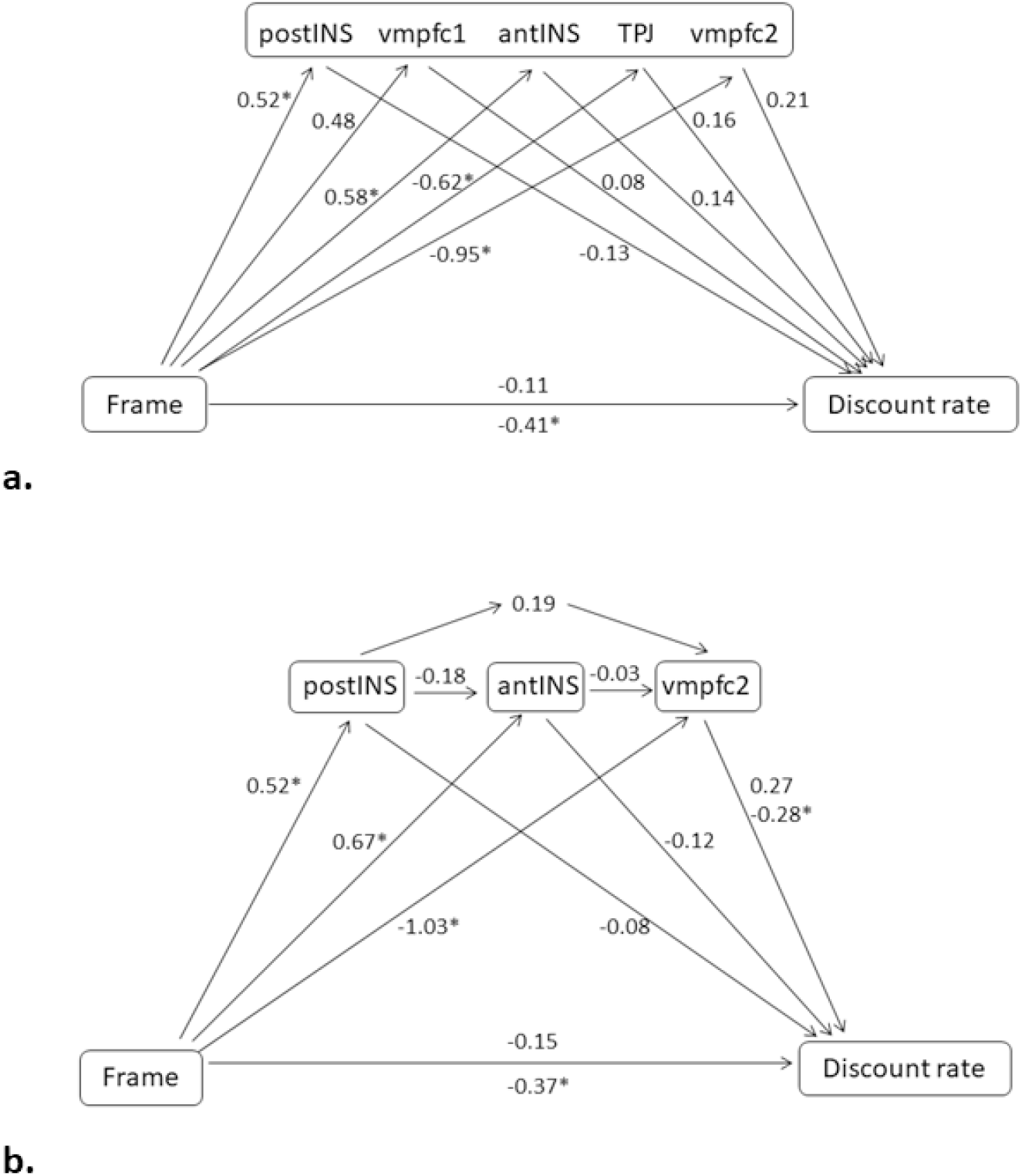
Mediation analyses. Two mediation models were built to clarify the effect of frame conditions (X) on the hyperbolic discount rate *k* (Y). Neural activations were included as mediators of such effect. In *model 1* (**a**), neural activations were entered as parallel possible mediators. In *model 2* (**b**), loss-specific, neural activations were entered as serial possible mediators. Numbers are partially standardized effects. *refers to significant effects (p < 0.05). Where two numbers are reported, the one on the top refers to the direct effect and the one on the bottom refers to the indirect effect.

To clarify this finding even more, we built, next, a loss-specific model (*model 2*) where mediators were treated serially. Based on the previous model, we included, in order, the posterior insula, the anterior insula, and VMPFC (GLM1) as moderators. While the direct effect of frame condition on the *k* parameter was not significant (p = 0.65), the total indirect effect was significant and negative, indicating that all mediators significantly mediated the relation between frame condition and *k*, and that the effect was stronger on the loss than the gain frame (partially standardized B = −0.37, SE = 0.18, 95% biased-corrected CI, −0.76 to −0.04). However, VMPFC seemed to play a crucial role as it mediated significantly the influence of frame on *k*, and this mediation was significantly stronger than in combination with insula activation as well as than insula activation alone (all partially standardized Bs < −0.31, all SEs < 0.17) (**Fig. 8b**).

In conclusion, our results suggest that the frame effect on social discounting was mediated by the interplay between insula and VMPFC in the loss frame, and between TPJ and VMPFC in the gain frame. Thus, we maintain that the most parsimonious explanation of insula activation and its negative modulatory interplay with VMPFC is indeed a frame-dependent downregulation of own-reward values in the valuation network during social discounting, thus biasing participants away from selfish toward generous choices, while TPJ-VMPFC coupling in the gain frame reflects the upregulation of vicarious reward value signals in VMPFC, hence promoting altruism by increasing the attractiveness of the generous option. Thus, in brief, the different motives underlying generosity in the gain and the loss frame are reflected by differential, frame-dependent activation and connectivity patterns in the brain.

## Discussion

We provide behavioral and neural evidence for a simple nudge that aims at increasing individuals’ willingness to provide costly support to socially remote others. We adapted a social discounting task where participants chose between a selfish option – a high gain to self and zero-gain to the other – and a generous option – a lower gain to self and equal gain to the other (Soutschek et al., 2016; Strombach et al., 2015). Based on previous evidence that people are strongly reluctant to increase their own payoff at the expense of others’ welfare (Baumeister et al., 1994; Chang & Sanfey, 2013; Chang et al., 2011; Crockett et al., 2014), we framed the generous option either as a monetary gain to the other (gain frame), or as the prevention of the loss of a previous monetary endowment to the other (loss frame) (Everett et al., 2015; Li et al., 2017; List, 2007; Liu et al., 2020; Sip et al., 2015; Smith et al., 2015; Wang et al., 2016; Xiao et al., 2016; Zheng et al., 2010). Crucially, between frames, the choice alternatives differed only in the description of the decision problem, but not with regard to their actual economic consequences. In a series of four independent studies, we show that the social discount function was significantly flatter in the loss than the gain frame, indicating that participants were more generous towards socially remote others if a personal gain implied the other’s loss of their previous endowment. Notably, our incentivization procedure made it logically impossible for the other person to know about her endowment, or the potential loss of it, and participants were explicitly instructed about this; all that mattered was the final positive payoff to self and others. Yet, the fact that our participants were still reluctant to inflict losses to others suggests that they had internalized the social norm of not taking away money from others to such a degree that it prevailed even in the absence of any real economic consequences for others.

We hypothesized that the frame-dependent motives underlying generosity are dissociable on the neural level. Consistent with our previous work (Strombach et al., 2015), we found that generosity in the gain frame recruited a network of structures, including VMPFC and rTPJ, known to represent vicarious reward value and overcoming egoism bias. By contrast, in the loss frame, we expected that the reluctance to maximize own-gain at the expense of other-loss would be mediated by social norm compliance and associated social sentiments, such as the negative emotions experienced during social norm transgressions, e.g., guilt and shame, as well as the aversive experience of unfairness and inequality. We therefore hypothesized that increased activity in brain regions associated with such social sentiments, specifically the insular cortex, would be associated with generous choices in the loss frame specifically (Bellucci, Feng, Camilleri, Eickhoff, & Krueger, 2018; Canessa et al., 2017, 2013; Civai et al., 2012; Corradi-Dell’Acqua et al., 2013; Huggins et al., 2018; Lallement et al., 2013; Lamm, Decety, & Singer, 2011; Montague et al., 2007; Oldham et al., 2018; Paulus et al., 2003; Samanez-larkin et al., 2008; Siebenthal et al., 2017; Singer et al., 2006; Sokol-hessner, Camerer, & Phelps, 2013; Sokol-hessner & Rutledge, 2019; Spitzer et al., 2007; Tomasino et al., 2013; Wagner, N’Diaye, Ethofer, & Vuilleumier, 2011; Wang et al., 2016; Yu et al., 2014). We indeed found that the anterior insula was significantly more activated when participants made generous choices in the loss frame, relative to the gain frame, and our analysis further showed that the degree of insula activation across trials correlated with the individual propensity to make generous choices in the loss, but not the gain frame. Extending these findings, we found that also the posterior part of the insula seemed to be involved in these processes, specifically supporting the representation of the loss frame information even before the decision was made (see also Droutman et al. 2015). Building upon this evidence, we further explored how both activation clusters mediated frame-specific social discounting behavior. We propose and provide empirical support for a network model that predicts that the frame effect on social discounting was associated with a frame-dependent neural connectivity pattern between insula and VMPFC in the loss frame, and TPJ and VMPFC in the gain frame. More specifically, DCM and mediation analyses confirmed that posterior insula activation at loss frame onset exerted a negative modulatory effect on VMPFC. This finding is consistent with the idea that a frame-dependent downregulation of own-reward values in the valuation network during social discounting might lie at the core of the enhanced generosity observed in the loss frame by biasing participants away from selfish toward generous choices. By contrast, the same analyses confirmed TPJ-VMPFC coupling in the gain frame, consistent with our previous finding (Strombach et al., 2015) that altruism in the gain frame is promoted by increasing the attractiveness of the generous option through TPJ-related upregulation of vicarious reward value signals in the valuation network. Overall, these results call for the idea that the motives behind generosity are likely qualitatively different in the gain and the loss frame, and dissociable on the neural level.

Our analyses revealed two separate clusters within insula; while a more posterior cluster was activated in response to general loss frame information, the more anterior cluster was specific to generous choices in the loss frame. This topographic dissociation within insula is consistent with previous findings suggesting a regional gradient in representing the level of abstraction of social sentiments during moral decision-making (e.g. Droutman et al. 2015; Ying et al. 2018). Yet, our analyses also showed that activation in both insula clusters correlated with the individual proportion of generous choices, and VMPFC connectivity with both insula clusters mediated the frame effect on social discounting. This pattern of result is in line with the idea that anterior and posterior insula may not subserve qualitatively different functions, but rather different aspects of the same function, such as the interoceptive and visceral aspects of social sentiments in response to potential vicarious feelings of potential loss (posterior cluster), and their relevance for choice selection (anterior cluster). Future research needs to clarify the specific functional differentiation of anterior versus posterior insula contributions to social economic decision-making.

Our findings expand on previous evidence that preventing harm to others is a great motivator of prosocial performance (Everett et al., 2015; Wang et al., 2016; Xiao et al., 2016; Zhang, Liu, Chen, Shang, & Liu, 2017; Zheng et al., 2010). However, while others have found that harm prevention was particularly pronounced in a public context (Everett et al., 2015) and dependent on social feedback (Sip et al., 2015; Smith et al., 2015), we show that similar cognitive mechanisms can strongly boost generosity even in a private context and in the absence of social feedback, thus independent of reputational concerns, judgment by social peers, or third-party punishment threats. This suggests that other-harm prevention might be an internalized motive that works unconditionally and universally across contexts, regardless of social consequences. In addition, previous experiments on harm prevention did not manipulate, or provide information on, social distance between donor and recipient (Bardsley, 2008; Crockett et al., 2014; Everett et al., 2015; Li et al., 2017; Liu et al., 2020; Xiao et al., 2016). Hence, while the effects of the resource allocation mode on social discounting were elusive so far, our findings imply that it matters: harm-prevention motives in the loss frame were less dependent on social distance than other-regarding considerations in the gain frame, thus resulting in flatter social discounting.

A recent study used a similar framing manipulation and also reported TPJ involvement (Liu et al., 2020). However, their study differed from ours in several important ways. First, the task in Liu et al. (2020) involved trading off own-wealth maximization with avoiding electric shocks to others. Thus, their task did not involve social distance information about the recipients of shocks. Second, Liu et al. (2020) did not reveal any insula recruitment, or insula-VMPFC connectivity - the core finding in our study - related to generosity or task framing. Most importantly, perhaps, while Liu et al. (2020) identified TPJ-VMPFC connectivity to be relevant for their frame-related increase in costly harm-prevention, we found instead that insula-VMPFC connectivity was associated with the frame-related boost in generosity during social discounting. This suggests that Liu et al. (2020) most likely studied different framing-related mental and neural mechanisms than the ones investigated here.

Our results are consistent with the idea that certain costly altruistic behaviors are not motivated by genuinely other-regarding considerations, but instead by compliance to internalized social norms. But what impels participants to comply to social norms? Here, we propose, along with previous evidence (Chang et al., 2011; Spitzer et al., 2007), that compliance to social norms might be linked to anticipated feelings of guilt, shame, and remorse, and accompanied by insula activation (see also Belfi et al. 2015; Sellitto et al. 2016), which ultimately sustain prosocial behavior.

The acceptance and support of the principle of a caring society, and the attitude towards the welfare of socially remote strangers, is central for a civilization to function well. It seems vital for societies to successfully meet current challenges, such as integrating refugees, addressing economic inequality, acceding the trials and promises of a globalized world, or managing the public health implications of the current COVID-19 pandemic (Kalenscher, 2014). Here, we present a simple behavioral framing manipulation that boosts generosity towards socially remote others: framing a selfish choice as a loss to others can result in a negative affective state, eventually motivating prosocial behavior, even if the framing of the choice options is irrelevant for the actual payoff to others. Our neuroimaging data identify insula as the core component in processing loss prevention motives, prompting generosity by biasing people away from selfish desires if these caused harm to others, including strangers. Our results imply that prosocial attitudes towards others are highly malleable and strongly depend on the architecture of the decision problem. The insights gained in this study might, thus, help in designing policies aimed at increasing the acceptance and support of the principle of a caring society, and to change the attitude towards the welfare of socially remote strangers.

## Materials and methods

### Participants

#### Studies 1-3

Three separate behavioral studies were carried out to test the validity of our paradigm in different settings and with different compensation procedures. For these studies we did not calculate the sample size in advance as we were not aware of any previous similar manipulation of social discounting. Study 1 was run online (*n* = 61; nine participants later excluded from the hyperbolic model fitting due to bad fitting; 27 females; mean age = 36 years, ±10.5 standard deviation) and participants were paid a fixed allowance of €8.5. Study 2 was run online (*n* = 36; six participants later excluded from the hyperbolic model fitting; 27 females; mean age = 21 years, ±2.7) and participants, all psychology students on campus, were reimbursed for their time with a fixed amount of university credits. Study 3 (*n* = 39; eight participants later excluded from the hyperbolic model fitting; 24 females; mean age = 25 years, ±5.7) was run in the laboratory and participants were paid out with the same fully incentive-compatible procedure as in the fMRI study 4 (see below). All three studies were conducted according to the Declaration of Helsinki and they were approved by the local ethics review board of the Heinrich-Heine University Düsseldorf. For studies 1 and 2 we did not collect informed consent, as this is allowed by the local ethics committee for online studies, which were completely anonymized, whereas we collected written informed consent in study 3, in the laboratory.

#### Study 4

After having replicated our results across the three behavioral studies with a within-subject design (see Results), for the fMRI study we estimated, via G*Power, assuming a medium-to-large effect size, that the sample size necessary to achieve a power of 0.95 was *n* = 23. Considering frequent participants’ drop out during long scanning sessions as our, or due to excessive movement, we opted for *n* = 40. Forty healthy young volunteers were therefore recruited at the Life&Brain Research Center in Bonn for an fMRI study. All participants met MR-compatible inclusion criteria in addition to no self-reported current or history of neurological or psychiatric disorder, as well as no current use of medication affecting the central nervous system. Due to excessive head motion during measurements (>4mm, >4° rotation, as computed through Artrepair Toolbox; Stanford Psychiatric Neuroimaging Laboratory, see (Cho et al., 2013; Strombach et al., 2015; Wendelken, O’Hare, Whitaker, Ferrer, & Bunge, 2011), 10 participants were excluded from all analyses. Thus, the final sample included 30 subjects (21 females; mean age = 25 years, ±4.6, range: 19-35 years) with high-education level (mean education = 14 years, ±1.9, range: 12-18 years, from high school to university master degree). Fifteen participants had a net monthly income between €0 and €499, eight between €500 and €999, five between €1000 and €1499, one between €1599 and €1999, and one larger than €2500.

All participants were fluent German speakers, right-handed, and had normal or corrected-to-normal vision. As reimbursement, they were paid €20 as participation fee, plus earnings from the social discounting task. Therefore, participants’ payoff ranged from €27.5 up to €35.5 (see Social discounting task).

The study was conducted according to the Declaration of Helsinki and it was approved by the local ethics review board of the Universitätsklinikum Bonn. All volunteers gave written informed consent to participate in the study.

### Social discounting task

In this task (adapted from Strombach et al. 2015), participants were first asked to imagine people from their social environment represented on a scale ranging from 1 (the person socially closest to them) to 100 (a random stranger), where a person at rank 50 was described as a person that the subject had seen several times without knowing the name. They were instructed to select six real persons located at social distances of 1, 5, 10, 20, 50, and 100 (with no need of specifying the name and their social relationship for the social distances 50 and 100). Participants were encouraged to avoid thinking of people that they felt negatively toward and people they shared a bank account or household with. Each trial began with the display of the social distance level of the partner the participant was playing with. Social distance was represented with a ruler scale consisting of 101 icons. The left-most icon, highlighted in purple, depicted the participant. One of the remaining 100 other icons was highlighted in yellow, indicating the social distance of the partner. Furthermore, social distance information was additionally indicated as a number on top of the highlighted yellow icon to prevent perceptual inaccuracies in estimating social distance (cf. **Fig. 1** for an example on a partner on social distance 100).

We included two experimental conditions, a *gain frame* and a *loss frame*. The *gain frame* manipulation was near-identical to the task used by (Strombach et al. 2015). Briefly, after presenting the social distance information as described above, participants were instructed that, in this trial, the experimenter gave an initial endowment of €0 the other person (“This person has €0”; **Fig. 1a**). Participants were explicitly and repeatedly instructed that the other person was not aware of her zero endowment, she would only be informed of the final payoff after implementing the participant’s choice. Then, two monetary options appeared, a selfish and a generous option. The selfish option (on the left in **Fig. 1a**, in purple letters) indicated the reward magnitude for the participant, if chosen (e.g., €115 to the participant and no other-reward to other). The generous option contained a smaller own-reward to the participant (€75) and an other-reward to the other person (€75). Own-rewards were always indicated in purple and other-rewards were always indicated in yellow. Participants indicated their choice of the selfish or the generous alternative by a left or right button press.

In the *loss frame* (**Fig. 1b**), participants were informed, after the social distance presentation, that the other person has received an initial endowment of €75 (“This person has €75”; **Fig. 1b**). As before, participants were explicitly and repeatedly instructed that the other person was not aware of her initial endowment, or the potential loss of it. On the next screen, a selfish (on the left in **Fig. 1b**) and a generous alternative (on the right in **Fig. 1b**) appeared. When choosing the selfish alternative, the participant received the own-reward amount indicated in purple (here, €115), and the other person lost her initial endowment, as indicated in yellow (-€75), thus leaving her empty-handed. When choosing the generous alternative, the participant received a smaller own-reward indicated in purple (€75), implying that the other person would keep her endowment.

In addition to the framing (gain frame, loss frame) and the social distance levels of the other (1, 5, 10, 20, 50, 100), in each condition, we manipulated the magnitude of the own-reward across trials: we used nine selfish reward amounts per frame condition, ranging from €75 to €155 in steps of €10. The generous alternative’s payoff was invariant, always yielding €75 own-reward and €75 other-reward in all conditions and trials.

Thus, in the gain frame condition, the other person always had a €0 endowment, the selfish alternative always yielded a variable own-reward, and no reward for the other, and the generous alternative invariantly yielded an equal €75/€75 split between participant and other person. In the loss frame condition, the other-endowment was always €75, the selfish alternative yielded a variable own-reward accompanied by the loss of the €75 endowment to the other, and the generous alternative always yielded €75 own-reward and had no financial consequences for the other, i.e., she could keep her initial endowment of €75.

To summarize the logic of the task, both frames were mathematically equivalent, i.e., they yielded identical final payoff states to the participant and the other person (in the example in **Fig. 1**: both frames yield an own-reward gain of €115 to the participant and €0 gain to the other person after a selfish choice, or €75 own-reward and €75 other-reward after a generous choice). The only difference between conditions was that a €0 other-reward outcome was framed as a loss of the initial endowment in the *loss frame* vs. a null-gain in the *gain frame,* and a €75 other-reward was framed as keep-endowment in the *loss frame* vs. a €75 gain in the *gain frame*.

The order of frame conditions, selfish-reward presentations, as well as the left or right screen-position of the selfish and generous alternative were randomized and counterbalanced across trials. The task of studies 1-3 had a total of 108 self-paced trials. The task of study 4 had a total of 216 trials as each trial type was repeated twice to allow for full left/right position counterbalancing. For events duration of study 4, please refer to **Fig. 1**.

#### Incentivization procedure

In studies 3 and 4 the social discounting task was fully incentive-compatible. At the end of the session, one of the participant’s choices was randomly drawn and 10% of the own-reward amount was paid out, as well as 10% of the other-reward amount was paid out to the other person in that trial, either via cheque, for the other-persons indicated by the participants at social distance 1, 5, 10, 20, or in cash to a random person on site in the case of other-persons at social distance 50 and 100. Note that the recipients of other-reward were only notified in case of a positive payoff, but not in case of a zero payoff or in case a trial was randomly chosen that did not consider them; in addition, they were not informed beforehand about this experiment, and, thus, had no prior outcome expectations. Hence, our incentivization procedure made it logically impossible for the other persons to know about their endowment, or the loss of it.

### General procedure

#### Studies 1-3

All participants performed a social discounting task and, at the end, they completed a questionnaire assessing social desirability (see Supplementary materials and methods). Although the social discounting task of studies 1 and 2 was not incentivized, participants were strongly encouraged to think as if they were making decisions for real. In studies 1 and 2, participants were instructed about the social discounting task, and then, after answering comprehension questions, they assigned other persons (i.e., name and personal relationship with them) from their social environment to the social distances 1, 5, 10, and 20, completed the task (see Social discounting task), and finally filled out a questionnaire (see Supplementary materials and methods) through Unipark online survey software (Unipark questback). Participants were provided with a web link to do so, after being recruited via flyers and advertisements on social platforms. Monetary payment for study 1 was implemented via Clickworker (GmbH), whereas university credits reimbursement was carried out on campus for study 2. In study 3, after recruitment, participants were invited to the laboratory, they were instructed on the social discounting task along with the comprehension questions. They then completed the task, implemented in Matlab R2016a (MathWorks) and Cogent toolbox (www.vislab.ucl.ac.uk), and filled out a questionnaire on a laptop. Finally, they were reimbursed for participation contingent on their choices, identical to the incentivization procedure in study 4.

#### Study 4

Upon arrival, participants received instructions about the social discounting task and then, after applying comprehension questions to check for full understanding of the task, they assigned other persons (i.e., name and personal relationship with them) from their social environment to the social distances 1, 5, 10, and 20 via paper and pencil. Afterwards, participants performed a few sample trials to familiarize with the task structure and they were subsequently cleared for the scanning session. At the end of the scanning session, they answered control questions concerning the social discounting task, and filled out a demographic questionnaire as well as questionnaires assessing social desirability and empathy (see Supplementary materials and methods). Finally, participants were debriefed and received their monetary allowance.

### Behavioral data analysis

#### Hyperbolic model

Similar to previous studies (Jones & Rachlin, 2006; Margittai et al., 2015; Margittai et al., 2018; Soutschek et al., 2016; Strombach et al., 2015), we approximated the participants’ decay in generosity across social distance with a hyperbolic function:

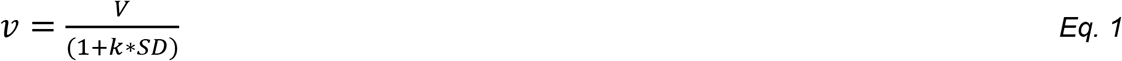

where v represents the discounted value of generosity, SD represents social distance, *k* represents the degree of discounting, and V is the intercept at social distance 0, thus the origin of the social discount function (Jones & Rachlin, 2006; Margittai et al., 2015; Soutschek et al., 2016; Strombach et al., 2015). While V can be considered an indicator of generosity towards socially close others (Margittai et al., 2015; Margittai et al., 2018; Strang et al., 2017), *k* describes the discount rate, i.e., the steepness by which the social discount function decays across social distance. We estimated *k* and V for each participant separately, depending on her individual choice pattern.

To estimate V, we titrated the selfish amount to determine, at each social distance, the point at which the subject was indifferent between the selfish and generous options (i.e., indifference point; see Supplementary results). Logistic regression, implemented in Matlab R2016a (MathWorks), was used to determine the indifference points where the likelihood of choosing the selfish and the generous options was 50% (Soutschek et al., 2016; Strombach et al., 2015).

To fit Eq. 1 and estimate *k*, we modeled trial-by-trial choices via a softmax function to compute the probability P of choosing the selected option o_i_ on a given trial:

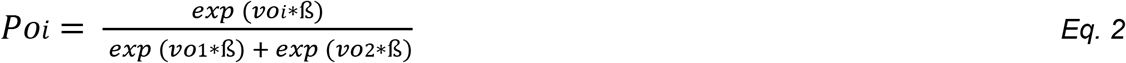

given the subjective values v (based on the current selfish amount and social distance) of the current available options o_1_ (vo_1_) and o_2_ (vo_2_) as in Eq. 1. ß is a nuisance parameter reflecting the stochasticity of individual performance. The larger ß, the less noisy the choice pattern. Individual discount rates were defined by the respective *k* value that yielded the best prediction of the observed choice probabilities by applying maximum-likelihood estimation using nonlinear optimization procedures (fminsearch function), implemented in Matlab R2016a (MathWorks). To this end, we minimized the log-likelihood of the choice probabilities to obtain the best-fitting *k* and ß parameter estimates, by summing across trials, given a specific set of model parameters *k* and ß, the logarithm of P(o_i_).

Both variables V and *k* were analyzed using non-parametric statistics as they were, in most of the cases across the four studies, not normally distributed (Kolmogorov-Smirnov, all ds > 0.20, all ps < 0.05). When participants did not discount at all (i.e., they always chose the generous option), *k* was set to 0 and V was set to 80 (i.e., maximum reward amount foregone = maximum selfish amount 155 – generous amount 75). All behavioral analyses were run in Statistica 12 (StatSoft). For additional analyses (i.e., indifference points, area under the curve, reaction times, and questionnaires) please see the Supplementary information.

### Study 4

#### fMRI procedures

Magnetic resonance images were collected on a 3T whole-body scanner (Magnetom Trio, Siemens Medical Systems, Erlangen, Germany) with an 8-channel head coil. For functional imaging, gradient-echo echo-planar images (EPI) were acquired at TR = 2500 ms (TE = 30 ms; number of slices = 37; slice thickness = 3 mm; distance factor = 10%; FoV = 192 mm × 192 mm; matrix size = 64 × 64; flip angle = 90°). Slices (voxel size = 2 × 2 × 3 mm) were sampled in transversal orientation covering all of the brain, including the midbrain. The scanning session started with a brief localizer acquisition. Afterwards, functional data were acquired in 3 separate runs of ~421 volumes each, to allow for brief resting periods in between. In order to get information for B_0_ distortion correction of the acquired EPI images, a gradient echo field map sequence (TR = 392 ms; TE 1 = 4.92 ms; TE 2 = 7.38 ms; number of slices = 37; voxel size 3 × 3 × 3 mm) was recorded before each functional run. Structural images were collected at the end (~5 min), using a T1-weighted sequence (rapid acquisition gradient echo sequence; 208 sagittal images; voxel size = 0.8 × 0.8 × 0.8 mm; 0.80 mm slice thickness). Head movements were minimized by the use of foam pads and scanner noise was reduced with earplugs. When necessary, vision was corrected-to-normal via fMRI compatible goggles. The social discounting task was programmed via an in-house software and presented via a mirror that projected a screen lying behind the participant, who made their choices via a left and a right button boxes.

#### fMRI preprocessing

Imaging data were preprocessed and analyzed with Statistical Parametric Mapping (SPM12, Wellcome Trust Centre for Neuroimaging, University College London, UK) implemented in Matlab R2016a (MathWorks). After checking raw data quality for each participant using SPM Check Reg function (Stanford Psychiatric Neuroimaging Laboratory), all images were preprocessed by reorienting them according to the EPI SPM template and coregistered to the fieldmap via FieldMap toolbox. After the functional images were realigned und unwarped to the middle volume and all volumes for participants’ motion correction by using phase correction, ArtRepair toolbox (Mazaika, Hoeft, Glover, & Reiss, n.d.) was run in order to identify bad volumes. Bad volumes of participants included in the final sample were not repaired. However, we modeled these bad volumes as regressors of no-interest in the statistical analyses (see fMRI analyses). Finally, functional and structural images were coregistered and the images were spatially normalized based on segmentation of the anatomical image with resampling to 2 × 2 × 2 mm, and spatially smoothed using a 6 mm FWHM Gaussian kernel. High-pass temporal filtering (using a filter width of 128 s) was also applied to the data.

#### fMRI analyses

At the first-level analysis, trial-related activity for each participant (*n* = 30) was modeled by delta functions convolved with a canonical hemodynamic response function to model the effects of interest, as well as six covariates capturing residual motion-related artifacts, and a temporal derivative for each regressor of interest to account for slice timing differences. For each participant, relevant contrasts were computed for each general linear model (GLM) (see below for details) and entered into second-level random effect analysis. The following variables were considered in the analyses: the loss frame condition; the gain frame condition; generous choices; selfish choices. Comparisons were run via one-way Analyses of Variance (ANOVAs), within subject, and via one-sample t-tests, where appropriate.

***GLM1*** searched for differences in BOLD activations between frame conditions during generous choices, where the onset of a generous choice was defined as the participant’s button press to choose the generous option after the monetary options had appeared on the screen (see **Fig. 1**). It included an unmodulated regressor of all generous choices made in the loss frame condition and an unmodulated regressor of all generous choices made in the gain frame condition. Additionally, the selfish amount magnitude (see Social discounting task) was included as trial-by-trial parametric modulator of all main regressors, separately. In the main manuscript, we additionally considered reward foregone as a parametric trial-by-trial regressor. Note that reward foregone is a linear transformation of, and thus collinear with, the selfish reward magnitude; neural activations identified by this parametric regressor might therefore reflect selfish amount or reward foregone (see main text). RTs were used as duration to account for differences between gain and loss frames (see Supplementary behavioral results). Additionally, missed trials were included as regressors of no-interest and modeled with duration = 5s, i.e., the maximum time allowed to respond.

Please note that at the level of choice, where the choice onset was defined as the participant’s button press after the release of the monetary options at each trial, a full model including separate regressors for both frames (gain and loss) and both types of choice (selfish and generous), as well as the trial-by-trial selfish amount as parametric regressor, was possible only for sixteen participants. This was due to participants who had to be excluded because they never, or only very rarely (not in all experimental runs) chose the selfish alternative in the loss frame. To address potential statistical power concerns associated with small sample size and to attend to potential selective sampling biases, in addition to generous choice being our main focus, we ran instead the above-mentioned model. Nevertheless, results of this full model (***GLMS1***), as well as of a model including only selfish choices (***GLMS2***), are reported in the Supplemental materials for completeness.

***GLM2*** tested for the effect of frame condition, and therefore included an unmodulated regressor of the onsets of the loss frame condition and an unmodulated regressor of the onsets of the gain frame condition. The frame onset was defined as the trial start (see **Fig. 1**). The social distance was included as trial-by-trial parametric modulator of the frame onsets, separately for the gain and the loss frames. A stick function was used as duration.

An additional ***GLM3*** was built with the sole purpose of extracting, at the individual level, beta estimates from two ROIs, one with seed in the anterior insula peak [42, 4, −4], and one with seed in the posterior insula peak [34, −16, 8], obtained from the previous GLM1 and GLM2, respectively, via MarsBaR region of interest toolbox for SPM12 (Matthew Brett, Jean-Luc Anton, Romain Valabregue, 2002). It included an unmodulated regressor of all choices (pooled across generous and selfish choices) made in the loss frame condition and an unmodulated regressor of all choices (pooled across generous and selfish choices) made in the gain frame condition. Choice onset was defined as the participant’s button press at each trial to choose either the selfish or the generous option after both monetary options had appeared on the screen (see **Fig. 1**). Pearson’s correlations were run between beta estimates for the gain and the loss frame, separately, with the percentage of generous choices out of the total amount of choices made in the respective frame.

All whole-brain level results as well as ROI-based (see below) results were initially thresholded at p < 0.001 (uncorrected), minimum cluster size = 5 voxels, and then corrected at the cluster level for multiple comparisons (p < 0.05, family-wise error rate [FWE]). Bad volume onsets (as measured via ArtRepair toolbox; (Mazaika et al., n.d.)), modeled with a stick function, were included as regressors of no-interest in all the above GLMs.

We additionally conducted, where relevant (see main text) ROI analyses for VMPFC, TPJ, and insular cortex by using anatomical bilateral masks from the Harvard-Oxford Atlas (Jenkinson, Beckmann, Behrens, Woolrich, & Smith, 2012), and the SPM Anatomical Automatic Labeling Toolbox, version 3 (Rolls, Huang, Lin, Feng, & Joliot, 2020), via SPM12 in Matlab R2016a.

Results are displayed in the figures overlaid on the averaged normalized structural images of the 30 participants included in the fMRI analyses. The probability maps of the SPM Anatomy Toolbox, version 3.0 (Eickhoff et al., 2007), and Neurosynth (http://neurosynth.org) were used for double checking region localization throughout GLMs.

##### Dynamic causal modeling (DCM)

We used DCM analysis as implemented in SPM12. This analysis focused on the interplay between insula and VMPFC and between TPJ and VMPFC, addressing both (i) regions endogenous connectivity and (ii) condition specific modulation of the regions (driving inputs) and their connections (modulatory inputs). We therefore constructed a hierarchical model with regressors defining both frame conditions activations against the total baseline activation. Thus, we entered in the DCM: a regressor of no-interest for baseline connectivity (‘all trials’, used to correct for global activation) including onsets of the screen presenting the framing information and the social distance, and the onsets of the screen presenting the monetary options, at all trials; a regressor (‘all loss trials’) including onsets of the screen presenting the framing information and the social distance, and the onsets of the screen presenting the monetary options, for the loss frame trials; a regressor (‘all gain trials’) including onsets of the screen presenting the framing information and the social distance, and the onsets of the screen presenting the monetary options, for the gain frame trials.

Subject-specific coordinates were guided by ROI-based group activation maxima in the three network regions from the univariate, group-level results (see Results section). Volumes of interest (VOI) spheres, with a radius of 6 mm, were built around the posterior insula (GLM2, [34, −16, 8]), rTPJ (GLM1, [50, −66, 36]), and VMPFC (GLM2, [2, 50, −8]). Note that we focused our DCM analysis on the posterior insula cluster only, as we were interested in a baseline frame activation; including the anterior insula cluster, specific for generous choice within the loss frame (see Results section), might have biased the results in favor of our hypotheses. Also note that we found TPJ in the choice-related analysis (GLM1) only: in our opinion, it was still preferable to opt for this experimentally driven ROI rather than including an ROI taken from the literature. Regional time series were extracted as the first eigenvariate of the three network regions for ‘all trials’ and mean-corrected for the effect of interest F-contrast at a liberal threshold of p = 0.1. This threshold was lowered for some participants until all regions could be detected (Zeidman, Jafarian, Corbin, et al., 2019; Zeidman, Jafarian, Seghier, et al., 2019). Based on our univariate results, we constructed bilinear models (**Fig. 7**) where the endogenous connectivity across the three regions was always assumed. We specified models with nodes reciprocally connected, where the gain and loss frame were allowed to modulate all connections (Li et al., 2015). The resulting 15 models were grouped in two families: A and B. In family A, both condition-specific driving inputs and condition-specific modulatory inputs were assumed. In family B, only condition-specific driving inputs were assumed.

*Family A* included eleven models (**Fig. 7**). In model *1*, we assumed that the gain frame condition had direct inputs on VMPFC and TPJ, and a modulatory input on their connections; the loss frame condition had direct inputs on VMPFC and insula, and a modulatory input on their connections. In model *2*, the gain frame had a driving input on VMPFC, and a modulatory input on its connectivity with TPJ; the loss frame had a driving input on VMPFC and a modulatory input on its connectivity with the insula. In model *3*, the gain frame had a driving input on TPJ and a modulatory input on its connectivity with VMPFC; the loss frame had a driving input on the insula and a modulatory input on its connectivity with VMPFC. In model *4*, the gain frame had a driving input on VMPFC and a modulatory input on its connectivity with TPJ; the loss frame had driving input on the insula and a modulatory input on its connectivity with VMPFC. In model *5*, the gain frame had a driving input on TPJ and a modulatory input on its connectivity with VMPFC; the loss frame had a driving input on VMPFC and a modulatory input on its connectivity with the insula. Therefore, connectivity between regions in model 1 to 5 is assumed to be bidirectional. Additionally, in model *6*, the gain frame had a driving input on VMPFC and a modulatory input on its connectivity to TPJ; the loss frame had a driving input on VMPFC and a modulatory input on its connectivity to the insula. In model *7*, the gain frame had a driving input on VMPFC and a modulatory input on the connectivity from TPJ to VMPFC; the loss frame had a driving input on VMPFC and a modulatory input on the connectivity from the insula to VMPFC. In model *8*, the gain frame had a driving input on TPJ and a modulatory input on its connectivity to VMPFC; the loss frame had a driving input on the insula and a modulatory input on its connectivity to VMPFC. In model *9*, the gain frame had a driving input on TPJ and a modulatory input on the connectivity from VMPFC to TPJ; the loss frame had a driving input on the insula and a modulatory input on the connectivity from VMPFC to the insula. In model *10*, the gain frame had a driving input on TPJ and a modulatory input on the connectivity from VMPFC to TPJ; the loss frame had a driving input on VMPFC and a modulatory input on its connectivity to the insula. In model *11*, the gain frame had a driving input on TPJ and a modulatory input on its connectivity to VMPFC; the loss frame had a driving input on VMPFC and a modulatory input on the connectivity from the insula to VMPFC.

*Family B* included four models (**Fig. 7**). In model *12*, the gain frame had driving input on VMPFC and TPJ; the loss frame had driving inputs on VMPFC and the insula. In model *13*, the gain frame had a driving input on TPJ and the loss frame had a driving input on insula. In model *14*, both frame conditions’ driving inputs were on VMPFC. In model *15*, the gain and the loss frame had driving inputs on all three regions, insula, TPJ, and VMPC, to check whether at increased number of connections, the model fits the data better.

All the hypothesized models were entered into Bayesian Model Selection (BMS), as implemented in SPM, to determine the best-fit family and model. The inference method used to compare the models across subjects and session was random effects (2^nd^-level, RFX). Bayesian Model Averaging (BMA) is used for model comparison. Once the optimal model was selected, the participant-specific parameters for the two frame conditions were averaged across the three runs and entered into group analysis with one-sample and paired-sample t-tests, where appropriate. This allowed us to summarize the consistent findings from the subject-specific DCMs using classical statistics (Cho et al., 2013; Li et al., 2015; Neufang et al., 2016; Wiehler, Petzschner, Stephan, & Peters, 2017; Zhang et al., 2018).

##### Mediation analyses

These analyses were run via Hayes’s PROCESS-macro (Hayes, 2017) as implemented in the IBM Statistical Package for the Social Sciences (SPSS). The analyses aimed at testing the idea that the gain and the loss frame had an effect on social discounting through the mediating influence of condition-specific neural activation. The frame condition was included as binary independent variable X (dummy variable: 1 = gain; 2 = loss), the hyperbolic *k* value (gain frame and loss frame) was entered as dependent variable Y, and the neural activations were entered as mediators. Specifically, beta estimates for the posterior insula [34, −16, 8; GLM2], VMPFC1 [2, 50, −8; GLM2], the anterior insula [42, 4, −4; GLM1], TPJ [50, −66, 36; GLM1], and VMPFC2 [0, 54, 14; GLM1] were extracted, at the single-subject level, for both frames and included in the model, via MarsBaR region of interest toolbox for SPM12 (Brett, Anton, Valabregue, 2002).

In *model 1* (**Fig. 8**), neural activations across both frames were treated as parallel mediators and included the posterior insula (GLM2) and the anterior insula (GLM1), TPJ (GLM1), and VMPFC (GLM1 and GLM2) (model template 4, *83*). In *model 2* (**Fig. 8**), loss-specific, mediators were treated serially (model template 6, *83*). Based on the previous model results, we included, in order, the posterior insula (GLM2), the anterior insula (GLM1), and VMPFC (GLM1) as moderators. Partially standardized values are reported, and 95% biased-corrected CIs are adopted. Number of bootstrap samples is set to 5000.

To determine the statistical power for mediation, the online tool MedPower was used (https://davidakenny.shinyapps.io/MedPower/) using effects of X on mediator (M) (path a), of M on Y (path b), and the direct effect of X on Y (path c′), at alpha=0.05. Total achieved power was ~0.60.

## Author contributions

M.S. and T.K. conceived the research project; M.S., A.S., B.W., and T.K. designed research; M.S. and A.S. performed research; M.S. analyzed data; M.S., S.N., and T.K. wrote the paper; and M.S., S.N., A.S., B.W., and T.K. revised versions of and approved the manuscript.

All authors declare no conflict of interest.

## Acknowledgments

This work was supported by Deutsche Forschungsgemeinschaft (DFG) grant no. KA 2675/4-3 (to T.K.).

## Data availability

Digital Imaging and Communications in Medicine (DICOM) images reported in this paper have been deposited in XNAT Central (https://central.xnat.org/) under the project name ‘Framesocdisc’. Behavioral datasets have been supplied in Figshare (https://figshare.com/) under the doi: 10.6084/m9.figshare.10265309.

## Supplementary information

### Supplementary materials and methods

#### Studies 1-3

##### Questionnaire

At the end of the social discounting task, participants completed the Social Desirability Scale (SDS-17; (Stöber, 2001)), a 16-item self-report questionnaire measuring the tendency to describe oneself with socially desirable attributes.

#### Study 4

##### Control questions

At the end of the scanning session, we asked participants to indicate, on a 7-point Likert scale (1 = not at all; 7 = definitely yes), their relationship with the persons they assigned to the various social distance levels during the social discounting task. These items included:

Item 1: “I feel very close to (person at social distance X)”;
Item 2: “In general, I have a very good relation with (person at social distance X)”;
Item 3: “I am very familiar with (person at social distance X)”.

##### Questionnaires

At the end of the scanning session, in addition to the SDS-17 (Stöber, 2001), participants completed the Interpersonal Reactivity Index (IRI; (Davis, 1983; C. Paulus, 2009)), a 16-item self-report questionnaire measuring the ability to empathize with others.

### Supplementary behavioral data analysis

#### Studies 1-4

##### Area under the curve (AUC)

We also characterized social discounting behavior using a model-free measure, the AUC of the empirically derived discount function, i.e., the curve plotting the indifference points against social distance (Sellitto et al., 2016). The larger the AUC values, the larger the preference for generous over selfish options.

### Supplementary fMRI data analysis

#### Study 4

##### GLMS1

This model (*n* = 16) looked at the choice onset, defined as the participant’s button press at each trial, after the monetary options appeared (see **Fig. 1**). It included separate unmodulated regressors for generous choices onsets in the loss frame, for selfish choices onsets in the loss frame, for generous choices onsets in the gain frame, and for selfish choices onsets in the gain frame. Additionally, the selfish amount magnitude was included as trial-by-trial parametric modulator of each regressor, separately. Reaction times (RTs) were used as duration to account for differences between gain and loss frames (see Supplementary behavioral results); missed trials were included as regressors of no-interest, with duration = 5s, which was the maximum reaction time allowed.

##### GLMS2

This model tested for the neural activations during selfish choices, where the onset of a selfish choice was defined as the participant’s button press to choose the selfish option after the monetary options had appeared on the screen (see **Fig. 1**). It therefore included an unmodulated regressor of all selfish choices, pooled across frame conditions, and the selfish amount magnitude (see Social discounting task) as trial-by-trial parametric modulator of the selfish choice onset. Reaction times (RTs) were used as duration to account for differences between gain and loss frames (see Supplementary behavioral results). Additionally, missed trials were included as regressors of no-interest and modeled with duration = 5s, i.e., the maximum time allowed to respond. Four participants could not be included in this analysis as they did not make any selfish choice in one or more runs of the task.

### Supplementary behavioral results

#### Study 1

##### Hyperbolic model

It is not possible to model the behavior of participants with null discounting, i.e., participants who always chose the generous option across all social distance levels. In those cases, we replaced the *k* value with 0 and the V value with 80 (for justification and reasoning, see Behavioral data analysis in the main text). However, this practice might have biased our results in favour of our hypothesis. We thus repeated the analyses on the differences in the *k* and V parameters between frame conditions, but this time treating those *k* and V parameters of participants with zero discounting as missing values. There was still a significant difference in the *k* parameter between frame conditions (median k_gain_ = 0.018 vs. median k_loss_ = 0.010; Z = 2.55, p < 0.011; r = 0.48) but no difference in the V parameter (median V_gain_ = 88 vs. median V_loss_ = 83; Z = 1.22, p = 0.22), supporting our conclusion that social discounting was flatter in the loss frame than the gain frame. The same pattern of results was found in studies 2-4.

#### Studies 1-3

##### Questionnaire

We pooled the SDS-17 scores across studies 1-3, and tested whether social desirability explained V nor *k* across frame conditions. However, the correlation did not reach significance (both ps > 0.25).

#### Studies 1-4

##### Indifference points

Our behavioral results can be seen even more clearly when comparing the indifference points between frame conditions –the selfish amount at which participant is indifferent between the selfish option and the generous option– at each social distance (see the Behavioral data analysis in the main manuscript for calculation details). In all four studies, indifference points were significantly higher in the loss compared to the gain frame for social distance 10, 20, 50, and 100 (Wilcoxon matched pairs test: all Zs > 2.61, all ps < 0.008, Bonferroni corrected), with the exception of social distance 10 in study 3 (p = 0.07), confirming the increased tendency in the loss frame to act generously toward socially remote individuals.

##### AUC

AUCs calculated for both frame conditions were significantly larger in the loss than in the gain frame across all four studies (Wilcoxon matched pairs test: all Zs > 4.16, all ps < 0.0001), further corroborating the conclusion of a frame-dependent difference in generosity.

#### Study 4

##### RTs

Mean RTs were normally distributed (Kolmogorov-Smirnov test, all ds < 0.15, all ps > 0.20). An ANOVA on RTs with frame (gain, loss) and choice (selfish, generous) as within subject factors yielded a significant effect of frame (F_1,25_ = 13.14, p < 0.002, η_p_^2^ = 0.34), a significant effect of choice (F_1,25_ = 15.58, p < 0.001, η_p_^2^ = 0.38), but no significant interaction frame × choice (F_1,25_ = 1.18, p = 0.29) (supplemental **Fig. S3**). Bonferroni-corrected post hoc comparisons showed that participants had slower responses in the loss frame compared to the gain frame (2064.5 ms vs. 1894.0 ms), and that they had slower RTs when making selfish compared to generous decisions (2117.4 ms vs. 1841.1 ms). These results indicate that it was generally more difficult to make a selfish choice, and that it was generally more difficult to make decisions in the loss than the gain frame (see (26) for a similar pattern of results). We considered RTs as events duration in our fMRI analyses to, thus, avoid potential choice-related RT confounds.

##### Control questions

We included a range of control questions in study 4 that probed the nature of the relationship of the participants to the respective people assigned to the various social distance levels. As social distance (SD) 1, ten participants indicated their mother/father, nine indicated their sibling, seven indicated their partner/husband/wife/fiancé(e), three indicated a close friend (non-sexual), and one defined the relationship to SD1 as ‘other’ (i.e., not included in the given options). We further asked participants to indicate their relationship with the person on SD 1 on a 7-point Likert scale (1 = not at all; 7 = definitely yes). Participants rated item 1 (“I feel very close to (person at SD1)”) with an average score of 6.0 (±1.9 standard deviation), item 2 (“In general, I have a very good relation with (person at SD1)”) with 6.3 (±1.7), and item 3 (“I am very familiar with (person at SD1)”) with 6.0 (±1.9).

As SD5, eighteen participants indicated a close friend (non-sexual), four indicated their mother/father, three indicated their sibling, four indicated their partner (two of which reported to be in an open relationship), and one indicated his/her uncle/aunt. Participants rated item 1 with a mean score of 5.7 (±1.8), item 2 with 6.2 (±1.5), and item 3 with 5.6 (±1.5).

As SD10, nine participants indicated a close friend (non-sexual), nine indicated their partner, two indicated their sibling, two indicated an acquaintance, two indicated their grandparent, two indicated a colleague, one indicated his/her roommate, and one indicated his/her father/mother. Two participants defined the relationship to SD10 as other. Participants rated item 1 with a mean score of 4.3 (±1.5), item 2 with 5.6 (±1.5), and item 3 with 5.0 (±1.7). As SD20, twelve participants indicated a colleague, six indicated their partner, five indicated an acquaintance, two indicated their mother/father, one participant indicated his/her cousin, one indicated a close friend (non-sexual), and one indicated his/her roommate. Two participants defined the relationship to SD20 as other. Participants rated on average item 1 with 3.6 (±1.5), item 2 with 5.3 (±1.3), and item 3 with 4.0 (±1.7).

Bonferroni-corrected comparisons revealed that the ratings of all three items significantly decreased from each social distance level to the next (all ps < 0.0001), with the exception of SD1 and SD5 (no item-score was significantly different; all ps > 0.13), and SD10 and SD20 (item 2 only reached trend level with p = 0.06). These results confirm that participants understood the social distance concept, followed the instructions of the social discounting task, and were sensitive to the differences between the SD levels.

##### Questionnaires

Social desirability (as measured via the SDS-17), and perspective taking, empathic concern, personal distress, and fantasy (as measured via the IRI) did neither correlate with the V nor with the *k* parameters in neither frame condition (all ps > 0.05).

### Supplementary fMRI results

#### Study 4

##### GLMS1

When contrasting generous and selfish choices within the loss frame, [generous choice > selfish choice] revealed a cluster of activation in the motor cortex (MNI peak coordinates at 16, −18, 62, p_FWE-corr_ < 0.01), whereas [selfish choice > generous choice] yielded clusters of activation mainly located in the ACC (−10, 48, 22, p_FWE-corr_ < 0.000) and in the precuneus (−6, −50, 10, p_FWE-corr_ = 0.02). The same contrasts within the gain frame did not yield any significant activation. When comparing choices between the loss and the gain frame, [generous choice, gain frame > generous choice, loss frame] revealed a cluster of activation in ACC (−10, 42, 2), whereas [selfish choice, gain frame > selfish choice, loss frame] yielded clusters of activation mainly located in the right putamen (28, 18, −2, p_FWE-corr_ = 0.03), left putamen (−26, 4, −2, p_FWE-corr_ = 0.005), and in the motor cortex (14, −28, 66, p_FWE-corr_ = 0.046) (supplemental **Table S3**). Additionally, the selfish amount magnitude, included as trial-by-trial regressor, did not parametrically modulate any of the above contrasts.

##### GLMS2

In this model we analyzed the neural activity during selfish choices in both frames and found that selfish choices, pooled across frame conditions, showed activity mainly in prefrontal (valuation) regions, including the anterior cingulate cortex (ACC) (MNI peak coordinates at 2, 36, 22, p_FWE-corr_ < 0.001) as well as the midbrain (−8, −12, −14, p_FWE-corr_ < 0.001) (supplemental **Table S4**). Additionally, selfish amount magnitude, included as trial-by-trial regressor, did not parametrically modulate neural activity in those brain regions (supplemental **Fig. S4**).

## Supplementary figures

**Figure S1.**
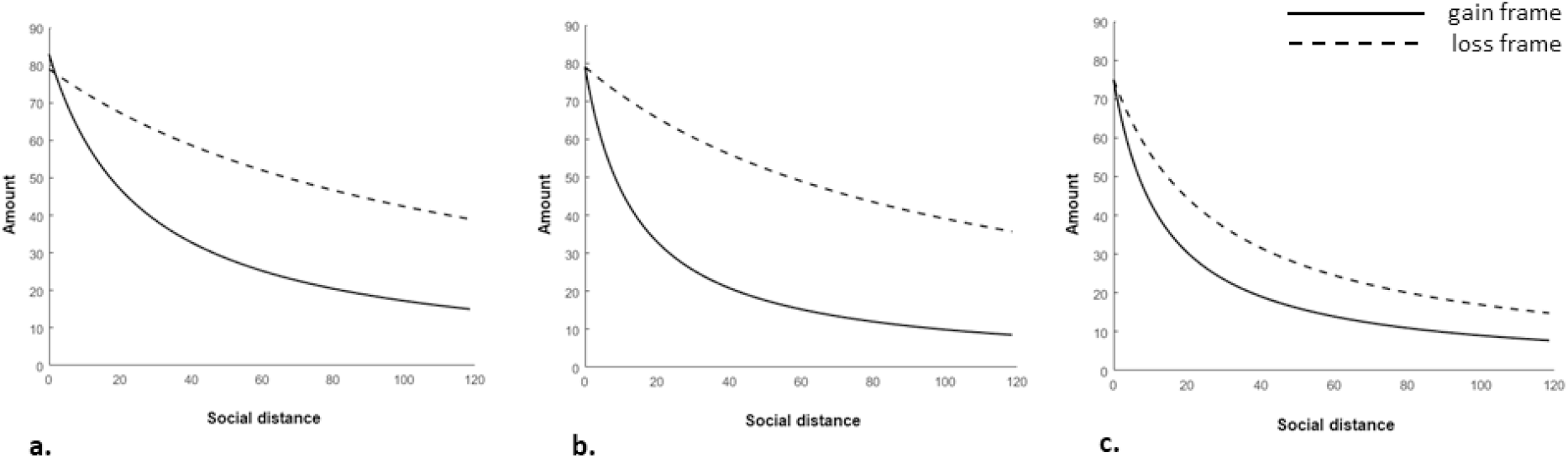
Hyperbolic discounting parameter *k*. The figure shows the mean best-fitting hyperbolic function across all participants of Study 1 (**a**), Study 2 (**b**), and Study 3 (**c**), separately for the gain frame (**solid line**) and the loss frame (**dashed line**). In all three studies, the social discount function was consistently flatter in the loss frame than in the gain frame.

**Figure S2.**
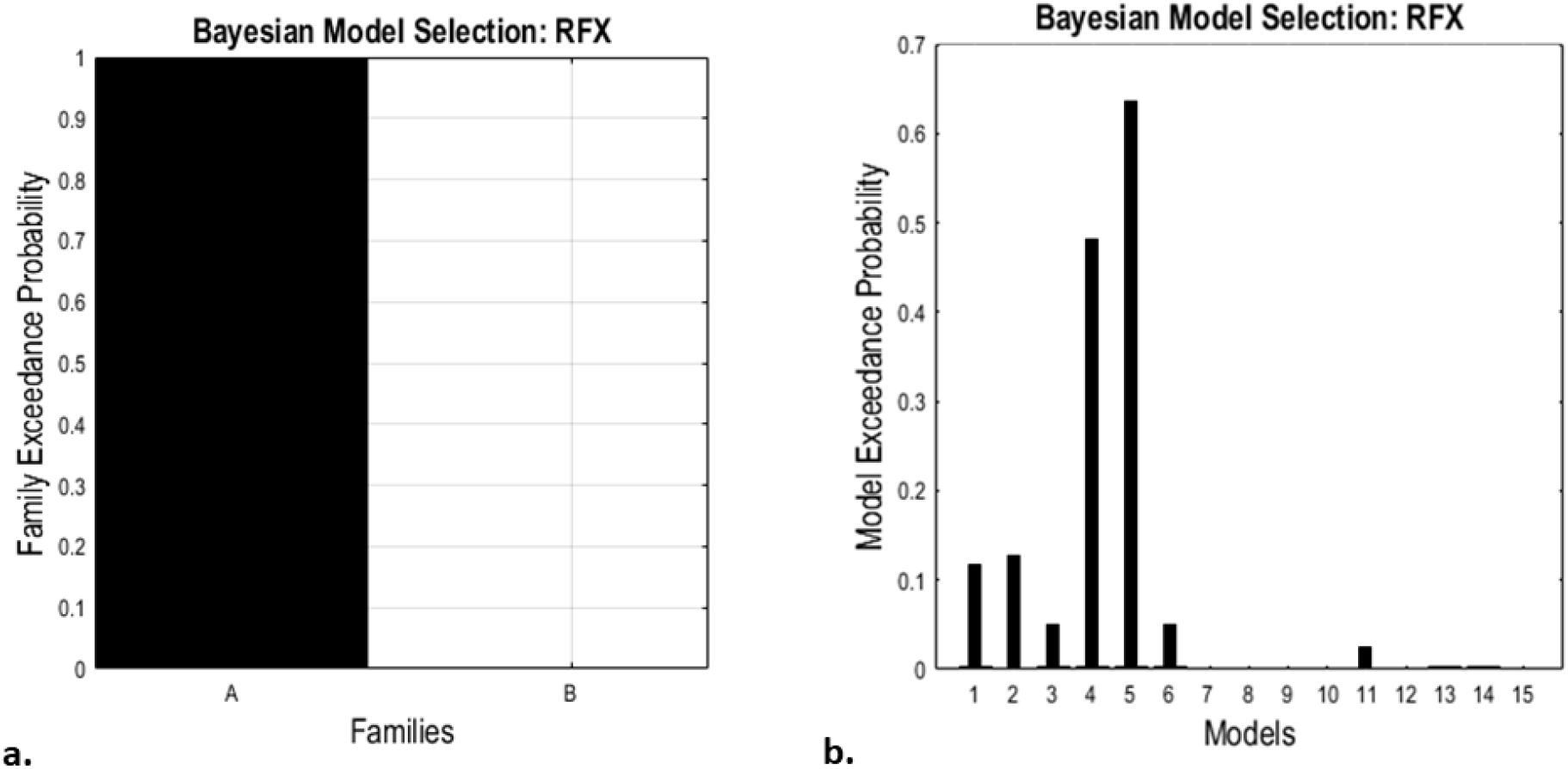
Bayesian model selection. Bayesian model selection revealed that the model family A (**a**), which assumed condition-specific effect at the level of both driving input and modulatory input, fits better the data, and that the model number 5 (**b**), which assumed that the gain frame had an effect on the TPJ and its connectivity with the VMPFC, while the loss frame had an effect on the VMPFC and its connectivity with the insula, is the winning one (sum of the log-evidence SF = −4.0786e+05, exceedance probability xp = 0.6508).

**Figure S3.**
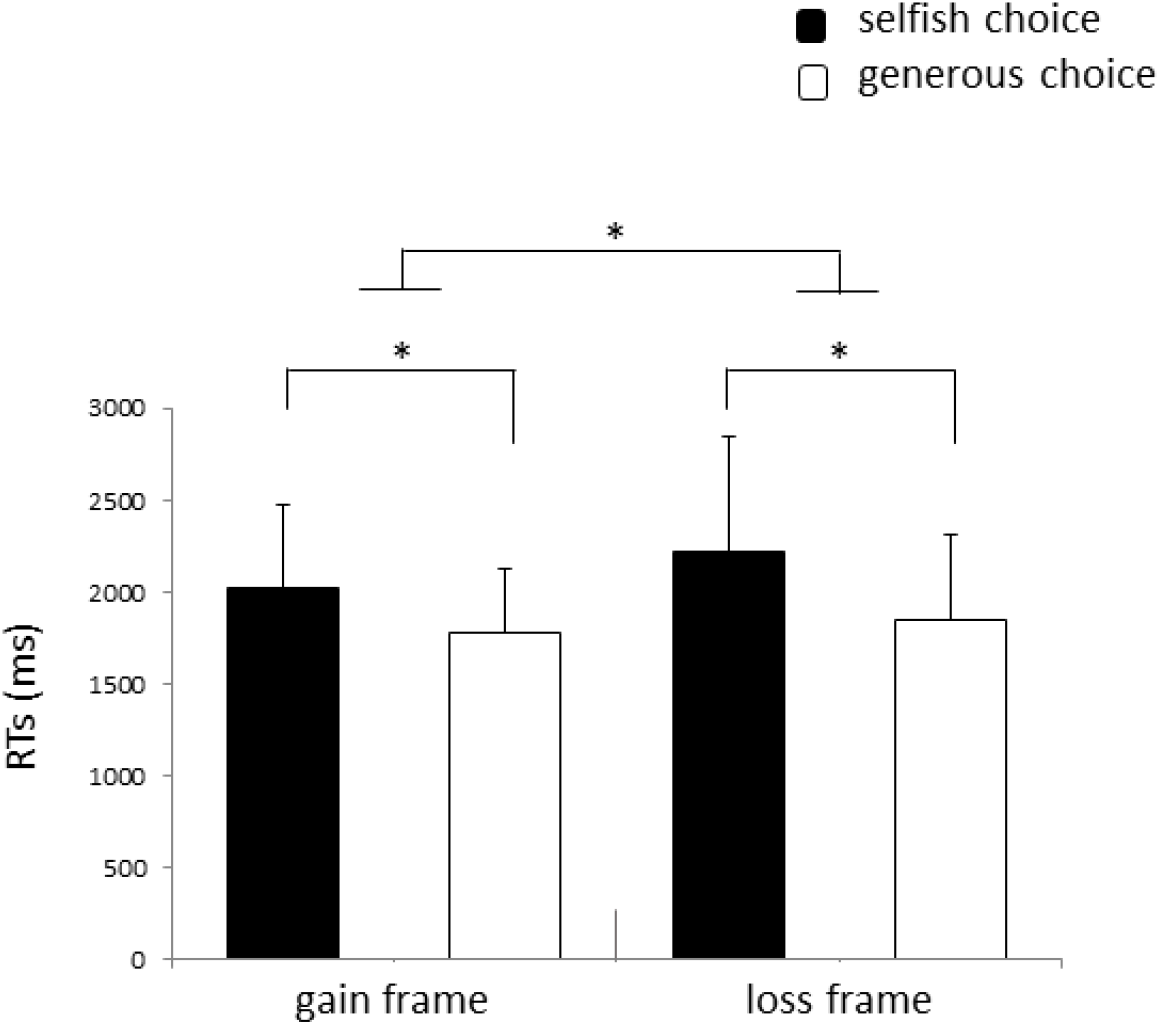
Reaction times. The figure shows the mean reaction times (RTs) expressed in milliseconds of Study 4. It took significantly longer to make a selfish than a generous choice, and it took significantly longer to make decisions in the loss than the gain frame. No interaction between choice type and frame was found. * indicates p < 0.005. Error bars indicate standard error.

**Figure S4.**
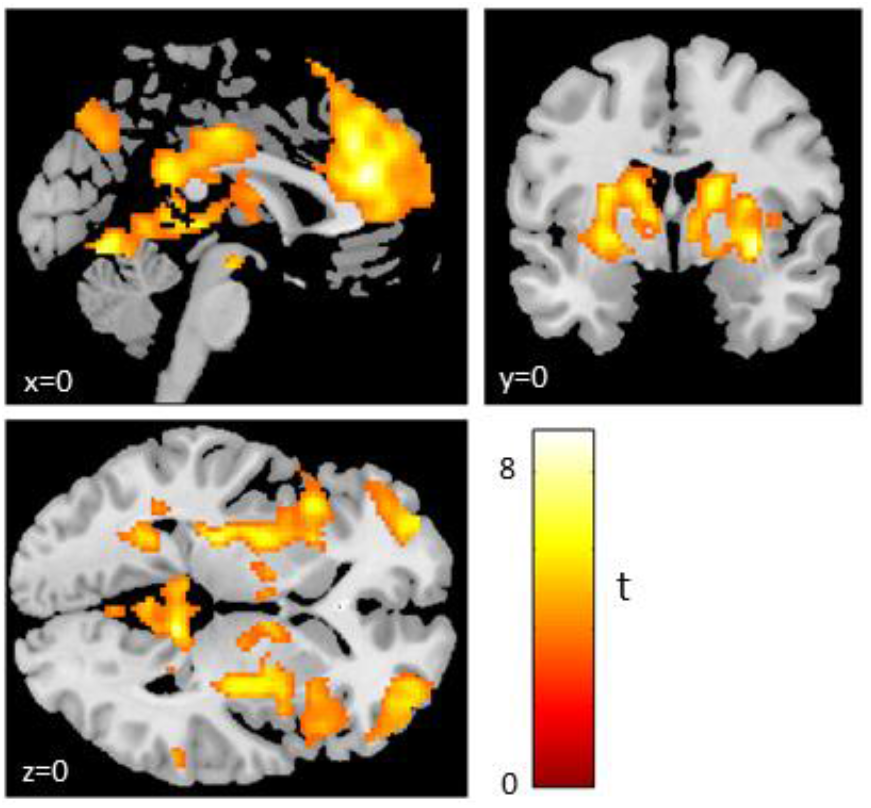
Selfish choices recruit the valuation network independent of frame. Selfish choices, pooled across gain and loss frames, activated the valuation network (GLMS1; p < 0.05 FWE whole-brain corrected at the cluster level; for illustration purposes, activations are displayed at p < 0.001, uncorrected, minimum cluster size ≥ 5). Color bar indicates T-value.

## Supplementary tables

**Table S1.** Activations at the generous choice onset (button press). Significant clusters active at p < 0.001 (unc.), whole-brain cluster-level-corrected p_FWE-corr_ < 0.05. L = left; R = right; ACC = anterior cingulate cortex; PCC = posterior cingulate cortex; VMPFC = ventromedial prefrontal cortex.

| **GLM1** |  |  |  |  |  |  | MNI coordinates |  |  |
| --- | --- | --- | --- | --- | --- | --- | --- | --- | --- |
|  | Region | Side | p<sub>FWE-corr</sub> | Cluster size | p(unc.) | Z-score | x | y | z |
| generous choice,<br>gain frame ><br>generous choice,<br>loss frame | VMPFC | L/R | 0.000 | 5424 | 0.000 | 6.59 | 0 | 54 | 14 |
|  | Frontal pole | L/R |  |  |  | 6.16 | 0 | 60 | 6 |
|  | ACC | L |  |  |  | 5.89 | -4 | 38 | 28 |
|  | PCC | L/R | 0.000 | 883 | 0.000 | 5.01 | 0 | -40 | 34 |
|  |  | L |  |  |  | 4.83 | -4 | -48 | 24 |
|  |  | L |  |  |  | 4.24 | -10 | -48 | 36 |
|  | TPJ | R | 0.034 | 124 | 0.002 | 4.65 | 50 | -66 | 36 |
|  | Parietal cortex | L | 0.021 | 139 | 0.001 | 4.35 | -44 | -74 | 36 |
|  |  | L |  |  |  | 3.59 | -44 | -64 | 26 |
|  | Caudate | L | 0.012 | 158 | 0.000 | 4.29 | -10 | 16 | 10 |
|  |  | L |  |  |  | 3.94 | -14 | 10 | 16 |
|  |  | L |  |  |  | 3.83 | -12 | 8 | -2 |
|  | Middle temporal<br>gyrus | L | 0.028 | 130 | 0.002 | 4.17 | -64 | -20 | -8 |
|  |  | L |  |  |  | 3.92 | -60 | -12 | -10 |
|  |  | L |  |  |  | 3.92 | -62 | -28 | -10 |
| generous choice,<br>loss frame ><br>generous choice,<br>gain frame | Precuneus | R | 0.000 | 284 | 0.000 | 4.05 | 8 | -54 | 54 |
|  |  | R |  |  |  | 3.89 | 10 | -62 | 62 |
|  |  | R |  |  |  | 3.63 | 12 | -70 | 58 |
|  |  | L | 0.004 | 194 | 0.000 | 3.94 | -8 | -66 | 58 |
|  |  | L |  |  |  | 3.81 | -8 | -54 | 64 |
|  |  | L |  |  |  | 3.72 | -12 | -50 | 54 |

**Table S2.** Activations at the frame and social distance information onset. Significant clusters active at p < 0.001 (unc.), whole-brain cluster-level-corrected p_FWE-corr_ < 0.05. L = left; R = right; ACC = anterior cingulate cortex; PCC = posterior cingulate cortex; VMPFC = ventromedial prefrontal cortex.

| *GLM2* |  |  |  |  |  |  | MNI coordinates |  |  |
| --- | --- | --- | --- | --- | --- | --- | --- | --- | --- |
|  | Region | Side | p<sub>FWE-corr</sub> | Cluster size | p(unc.) | Z-score | x | y | z |
| loss frame > gain frame | ACC | R | 0.000 | 302 | 0.000 | 5.03 | 4 | 32 | 22 |
|  |  | R |  |  |  | 4.58 | 4 | 22 | 16 |
|  | Caudate | R |  |  |  | 3.93 | 8 | 16 | 10 |
|  | Superior temporal gyrus | L | 0.004 | 181 | 0.000 | 4.96 | -48 | -14 | -2 |
|  | Middle temporal gyrus | L |  |  |  | 3.97 | -52 | -16 | -14 |
|  |  | L |  |  |  | 3.60 | -60 | -18 | -10 |
|  | PCC | L | 0.001 | 232 | 0.000 | 4.74 | -4 | -16 | 38 |
|  |  | L |  |  |  | 4.05 | -18 | -22 | 42 |
|  |  | L |  |  |  | 3.98 | -4 | -24 | 42 |
|  | Insula | R | 0.01 | 161 | 0.000 | 4.32 | 34 | -16 | 8 |
|  |  | R |  |  |  | 4.27 | 44 | -34 | 24 |
|  |  | R |  |  |  | 4.23 | 38 | -24 | 12 |
|  | Prefrontal lobe | R | 0.001 | 219 | 0.000 | 4.26 | 24 | 54 | 12 |
|  | VMPFC | L/R |  |  |  | 3.97 | 2 | 50 | -8 |
|  | ACC | R |  |  |  | 3.53 | 14 | 38 | -2 |
|  | Superior temporal gyrus | R | 0.005 | 170 | 0.000 | 4.18 | 58 | -16 | -10 |
|  |  | R |  |  |  | 3.82 | 52 | -2 | -16 |
|  | Premotor cortex | R |  |  |  | 3.27 | 50 | 4 | -22 |
|  |  | L |  |  |  | 3.52 | -18 | 22 | 38 |

**Table S3.**
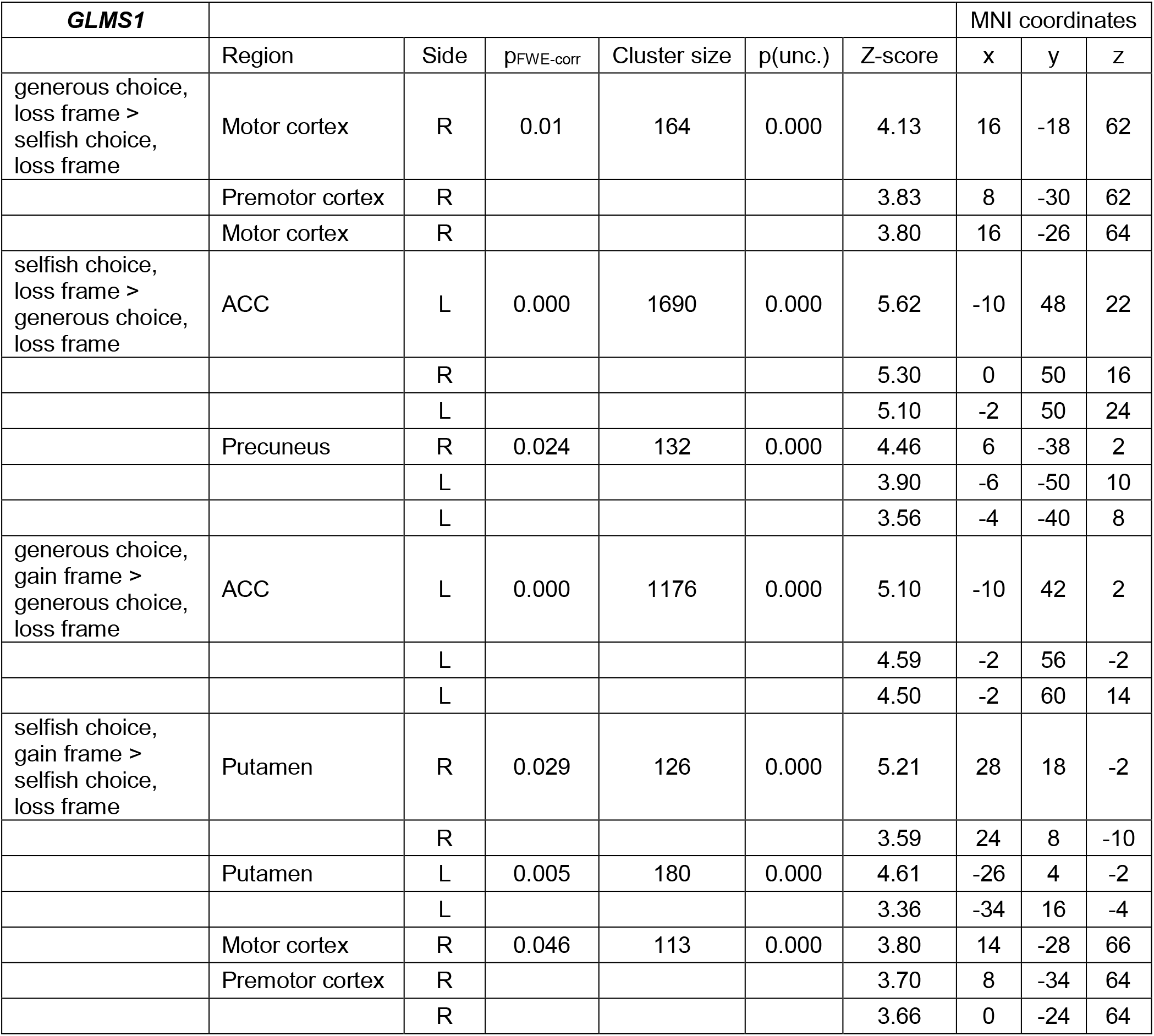
Activations at the choice onset (button press). Significant clusters active at p < 0.001 (unc.), whole-brain cluster-level-corrected p_FWE-corr_ < 0.05. L = left; R = right; ACC = anterior cingulate cortex.

**Table S4.** Activations at selfish choice onset (button press), pooled across loss and gain frame. Significant clusters active at p < 0.001 (unc.), whole-brain cluster-level-corrected p_FWE-corr_ < 0.05. L = left; R = right; ACC = anterior cingulate cortex.

| GLMS2 |  |  |  |  |  |  | MNI coordinates |  |  |
| --- | --- | --- | --- | --- | --- | --- | --- | --- | --- |
|  | Region | Side | p<sub>FWE-corr</sub> | Cluster size | p(unc.) | Z-score | x | y | z |
|  | Midbrain | L | 0.000 | 16664 | 0.000 | 5.96 | -8 | -12 | -14 |
|  | ACC | L/R |  |  |  | 5.87 | 2 | 36 | 22 |
|  | Thalamus | R |  |  |  | 5.80 | 22 | -24 | 12 |
|  | Parietal cortex | L | 0.000 | 1287 | 0.000 | 5.55 | -50 | -54 | 42 |
|  |  | L |  |  |  | 4.68 | -36 | -58 | 36 |
|  |  | L |  |  |  | 4.37 | -46 | -66 | 42 |
|  |  | R | 0.000 | 1762 | 0.000 | 5.41 | 50 | -54 | 50 |
|  |  | R |  |  |  | 5.11 | 54 | -56 | 42 |
|  |  | R |  |  |  | 5.09 | 46 | -50 | 42 |
|  | Frontal pole | R | 0.000 | 993 | 0.000 | 4.99 | 48 | 42 | -6 |
|  |  | R |  |  |  | 4.87 | 36 | 48 | 4 |
|  |  | R |  |  |  | 4.81 | 32 | 56 | 2 |
|  |  | L | 0.000 | 548 | 0.000 | 4.89 | -34 | 54 | -2 |
|  |  | L |  |  |  | 4.19 | -42 | 52 | 10 |
|  |  | L |  |  |  | 3.97 | -40 | 46 | 2 |
|  | Middle frontal gyrus | L | 0.003 | 203 | 0.000 | 4.74 | -44 | 18 | 48 |
|  |  | L |  |  |  | 4.12 | -40 | 14 | 42 |
|  | Precuneus | L/R | 0.000 | 449 | 0.000 | 4.14 | 2 | -78 | 46 |
|  |  | L/R |  |  |  | 4.12 | 0 | -68 | 36 |
|  |  | L |  |  |  | 3.87 | -6 | -62 | 36 |

## Notes

https://central.xnat.org/app/action/DisplayItemAction/search_value/Framesocdisc/search_element/xnat:projectData/search_field/xnat:projectData.ID

https://doi.org/10.6084/m9.figshare.10265309

### Competing Interest Statement

The authors have declared no competing interest.

### Summary of Updates

In this version of the manuscript mainly a DCM analysis has been included, thus providing additional findings.

